# β-Arrestin Condensates Regulate G Protein-Coupled Receptor Function

**DOI:** 10.1101/2025.04.05.647240

**Authors:** Preston J. Anderson, Peng Xiao, Yani Zhong, Adam Kaakati, Juliana Alfonso-DeSouza, Tianyao Zhang, Chao Zhang, Kevin Yu, Lei Qi, Wei Ding, Samuel Liu, Biswaranjan Pani, Athmika Krishnan, Oscar Chen, Chanpreet Jassal, Joseph Strawn, Jin-Peng Sun, Sudarshan Rajagopal

**Author notes:** These Authors equally contributed (Preston J. Anderson, Peng Xiao, Yani Zhong, Jin-Peng Sun, Sudarshan Rajagopal).

## Abstract

G protein-coupled receptors (GPCRs) are the largest class of receptors in the genome and control many signaling cascades essential for survival. GPCR signaling is regulated by β-arrestins, multifunctional adapter proteins that direct receptor desensitization, internalization, and signaling. While at many GPCRs, β-arrestins interact with a wide array of signaling effectors, it is unclear how β-arrestins promote such varied functions. Here we show that β-arrestins undergo liquid-liquid phase separation (LLPS) to form condensates that regulate GPCR function. We demonstrate that β-arrestin oligomerization occurs in proximity to the GPCR and regulates GPCR functions such as internalization and signaling. This model is supported by a cryoEM structure of the adhesion receptor ADGRE1 in a 2:2 complex with β-arrestin 1, with a β-arrestin orientation that can promote oligomerization. Our work provides a paradigm for β-arrestin condensates as regulators of GPCR function, with LLPS serving as an important promoter of signaling compartmentalization at GPCRs.

## INTRODUCTION

β-arrestins 1 and 2 are multifunctional adaptor proteins^1^ that regulate the signaling of G protein-coupled receptors (GPCRs), the largest class of receptors that impact nearly all aspects of physiology and are one of the most common drug targets^2^. After agonist binding, GPCRs activate heterotrimeric G proteins, which promote signaling through second messengers, followed by the recruitment of GPCR kinases (GRKs) and β-arrestins. Receptor phosphorylation by GRKs promotes associations with β-arrestins, wihch regulate GPCR desensitization, signaling, and trafficking. They perform these multiple tasks by sterically preventing G protein activation and by acting as scaffolding proteins for the endocytic machinery and signaling molecules (e.g., Raf-1, MEK1, and ERK) ^3,4^, interacting with hundreds of proteins^5^.

In addition, β-arrestins can be catalytically activated by GPCRs, followed by translocation to the plasma membrane (PM) and clathrin-coated pits (CCPs)^6^.

Structural studies of β-arrestins^7–9^ have demonstrated their interaction with the receptor through finger loop insertion into the receptor core and/or through binding of its N-domain to the phosphorylated receptor C-terminus^10–12^. This results in the release of the β-arrestin C-tail intrinsically disordered region (IDR) from the N-domain, and promotes the active confirmation of β-arrestin^13–15^ characterized by interdomain twisting between the N-and C-domains^16,17^. β-arrestins are regulated by their cellular environment through binding to specific components, such as the GPCR C-tail, inositols, and lipids^6,18–23^. In vitro studies have shown that inositol hexaphosphate (IP_6_) promotes β-arrestin 1 oligomerization into “infinite chains”^8^ and β-arrestin 2 oligomerization into trimers with a conformation similar to β-arrestin 1^12,24–26^ in complex with a GPCR ^8,27,28^. In cells, it has been shown that β-arrestins can homo-and hetero-oligomerize^29^, and ablation of inositol (IP_6_) binding sites results in dysregulation of β-arrestin nucleocytoplasmic shuttling, signaling, and protein:protein interactions^28,30^. However, the biological significance of β-arrestin oligomerization and whether it promotes compartmentalization of receptor signaling is unclear at this time.

One mechanism of compartmentalization that is known to contribute to signaling by other receptor families is the formation of biomolecular condensates^31,32^. Biomolecular condensates are enriched with molecules that can engage in multivalent interactions through specific oligomerization motifs and IDRs^33,34^. This decreases the solubility of the molecules due to entropy-driven effects and promotes liquid-liquid phase separation (LLPS) to produce condensates^31^. These condensates partition reactants to increase their local concentration and sequester signaling components^32,35,36^. Condensates have been described at membrane receptors such as receptor tyrosine kinases and T cell receptors,^36,37^ in addition to their downstream effectors, such as PKA and WNK^35,38,39^. Notably, β-arrestins 1 and 2 have a C-terminal IDR, and in the presence of IP_6,_ β-arrestin undergoes oligomerization^35–38,40–42^, features that are found in macromolecules enriched in biomolecular condensates.

Here, we demonstrate the formation of β-arrestin condensates to regulate GPCR activity. We demonstrate that mutations of the IDR and residues predicted to promote oligomerization alter the ability of β-arrestins to regulate GPCR signaling and internalization. Lastly, we determine the structure of the ADGRE1 receptor bound to β-arrestin 1 captured in a 2:2 stoichiometry in an engagement mode that can promote the formation of β-arrestin oligomers.

## RESULTS

### **β**-arrestins form condensates that are regulated by GPCR activation

To test for the ability of β-arrestins to oligomerize in vitro, we compared patterns of β-arrestin labeling with monomeric and split GFP variants. Labeling of β-arrestins with eGFP on the C-terminus of β-arrestin 1 or 2 demonstrated diffuse expression in the nucleus and cytoplasm for β-arrestin 1, and cytoplasm alone for β-arrestin 2 due to the presence of a nuclear export sequence in β-arrestin 2 **(Figure 1A and Supplemental Figure 1B)**^43^. To visualize potential interactions between β-arrestins, we coexpressed β-arrestin 2 labeled at its C-terminus with the 11^th^ β strand of split super folder GFP (sfGFP) (16 amino acids) along with β-arrestin 2 labeled at its C-terminus with the complementary sfGFP_1-10_ in HEK293 cells **(Figure 1B)**.

**Figure 1:**
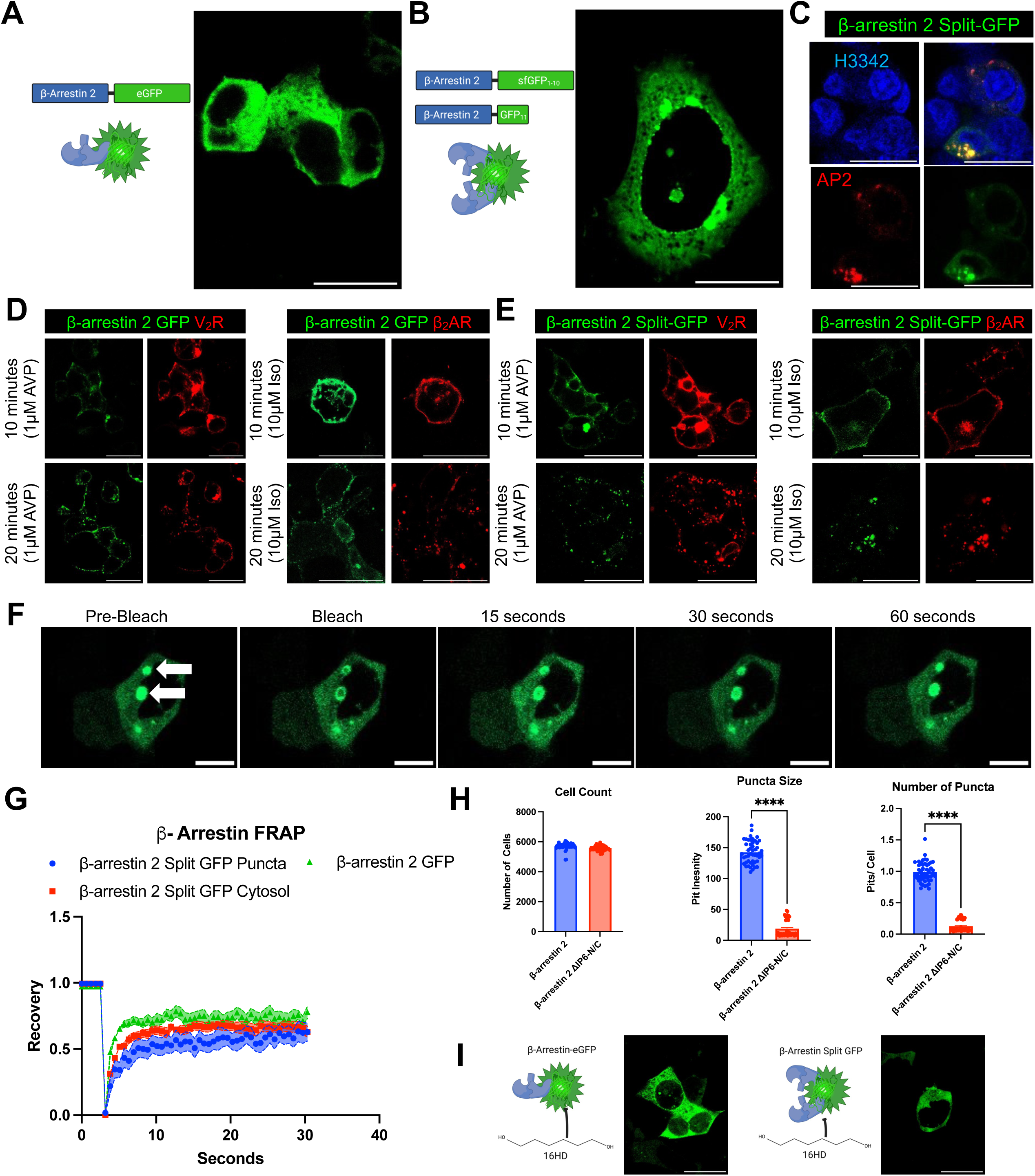
β**-arrestins form biomolecular condensates.** (**A**) Diffuse cytosolic β-arrestin 2-eGFP expression in HEK293 cells transfected with a β-arrestin 2 with eGFP fused to the C-terminus. (**B**) Observation of puncta β-arrestin 2 split-GFP system. β-arrestin 2 was fused with 11^th^ β-strand of GFP, or the remaining β-strands of the super folder GFP (sfGFP_10_) on the C-terminus. (**C**) Visualization of β-arrestin 2 puncta at clathrin-coated pits. Cells were transfected with β-arrestin-2 split-GFP, AP2-mKO (β subunit), and were stained with DAPI. (**D**) β-arrestin 2-eGFP recruitment upon agonist stimulation. Cells were transfected with β-arrestin-2-eGFP, and a fluorescently labeled receptor was stimulated with AVP (1 µM) or Isoproterenol (10 µM). Images were captured at 10 and 20 minutes. (**E**) β-arrestin-2 split-GFP recruitment upon agonist stimulation. Similar to the experimental design except cells were transfected with the β-arrestin-2 split-GFP system. (**F-G**) Monitoring the dynamics of cytosolic and β-arrestin 2 condensates. FRAP was performed on β-arrestin 2 split-GFP puncta and compared to cytosolic split-GFP and β-arrestin 2 eGFP. Curves show normalized average fluorescent intensity with SEM. (**H**) Quantification of size and number of puncta with of β-arrestin 2 split-GFP and IP_6_ mutants. (**I**) Left Panel: β-arrestin 2-eGFP exposure with 2.5% 1,6-Hexanediol (16HD) for 5 minutes. Right Panel: Same experimental design, except cells were transfected with the β-arrestin 2 split-GFP (H) Scale bars are 25µm for all images. Confocal microscopy images are representative of n=3.

Expression of β-arrestin GFP_11_ or β-arrestin sfGFP_1-10_ alone resulted in no observable signal **(Supplemental Figure 1A)**. With coexpression of both labeled β-arrestins, we observed the formation of puncta **(Figure 1B)**, consistent with an interaction between β-arrestins resulting in complementation of GFP_11_. These puncta localized with AP2, a marker of CCPs **(Figure 1C)**. Notably, we did not observe visible puncta with overexpression of β-arrestin 1 and 2-eGFP **(Supplemental Figure 1B)**. Similarly, titration of split-GFP labeled β-arrestin 1s resulted in puncta at low levels of overexpression (**Supplemental Figure 1C)**. Titration of both fluorescent β-arrestin systems (i.e., eGFP and split-GFP) was titrated near the endogenous expression of β-arrestin **(Supplemental Figure 1D)**.

To determine whether agonist activation of GPCRs had any effect on the signal generated by split-GFP β-arrestins, we overexpressed the β2-adrenergic (β_2_AR) or vasopressin receptor 2 (V_2_R) along with the labeled β-arrestins in HEK293 cells. These receptors display distinct patterns of β-arrestin recruitment to the receptor after agonist stimulation. The β_2_AR promotes a “class A” pattern, characterized by PM β-arrestin redistribution but minimal internalization with the receptor to endosomes while the V_2_R promoting a “class B” pattern, characterized by initial β-arrestin redistribution to the PM followed by strong localization to endosomes with the receptor.^44,45^ Agonist stimulation of the receptors transfected with β-arrestin 2-eGFP resulted in the expected patterns at 10 and 20 minutes **(Figure 1D)**. Cells transfected with β-arrestin split-GFP resulted in signal largely limited to the plasma membrane at both receptors at 10 minutes, similar to that observed with eGFP-labeled β-arrestins **(Figure 1E)**. In contrast, at 20 minutes split-GFP-labeled β-arrestins displayed distinct patterns from eGFP-labeled β-arrestins. While we observed some puncta at endosomes at the V_2_R, most of the signal of the split GFP after agonist stimulation persisted at the plasma membrane **(Figure 1E)**. This is consistent with distinct processes being monitored by eGFP and split GFP-labeled β-arrestin, namely redistribution and oligomerization, respectively. Together, these data are consistent with β-arrestin oligomerization being dynamically regulated by receptor activation.

To determine condensate fluidity, we next evaluated the dynamics of β-arrestin puncta with fluorescence recovery after photobleaching (FRAP) experiments **(Figure 1F-G and Video S1)**. Cytosolic β-arrestin 2-eGFP displayed nearly a ∼70% recovery (SEM± n=18) at 10 seconds. The β-arrestin 2 split-GFP system showed that puncta and diffusible cytosolic pools dynamically exchange, shown by similar recovery kinetics (cytosol SEM± n=15), although β-arrestin 2 split-GFP puncta had slightly slower kinetics (puncta SEM± n=13). We next tested if β-arrestin oligomerization regulates the size and number of these puncta. Using previously published β-arrestin IP_6_ oligomerization mutants^8^, termed β-arrestin ΔIP_6_-N/C, we quantified ∼5,000 cells and found a significant decrease in both puncta size and number **(Figure 1H).** We next tested the biophysical properties of these puncta by using 1,6 hexanediol (16HD), which disrupts pi-pi and hydrophobic interactions in condensates^40,46^. In cells overexpressing β-arrestin-eGFP, there was no change in the uniform distribution of β-arrestin-eGFP in the cytosol with 2.5% 16HD **(Figure 1I).** In contrast, in cells overexpressing β-arrestin split GFP, there was a loss of puncta formation with 2.5% 16HD **(Figure 1I)**. Together, these findings are consistent with β-arrestins forming condensates that are regulated by IP_6_-mediated oligomerization.

### Optogenetic control of **β**-arrestin LLPS

To dissect the components of β-arrestin needed for sufficient condensate formation, we leveraged a previously published optogenetic approach to modulate condensate formation^47^. In short, Cry2, upon exposure to blue light, induces oligomerization, and when fused with an intrinsically disordered region (IDR), drives LLPS to form condensates **(Figure 2A).** Consistent with previous reports, we found Cry2 WT (Cry2-mCherry) alone was not sufficient in driving condensate formation **(Figure 2B and Video S2)**^48,49^. Fusion with FUSN (N-terminus of FUS), a well-characterized IDR^50^ to Cry2WT, led to light-induced clustering in both the cytosol and the nucleus **(Figure 2C and Video S3)**. We next fused full-length β-arrestin 1 or 2, which we termed Opto-β-arrestins. For Opto-β-arrestin 1, light treatment promoted condensates formation within a matter of seconds, predominately in the cytosol **(Figure 2D and Video S4)**. In contrast, Opto-β-arrestin 2 displayed a few puncta at baseline, with minimal impact from light activation **(Figure 2E and Video S5)**. Together, this suggests that β-arrestin 1 and 2 may have different propensities to form condensates.

**Figure 2:**
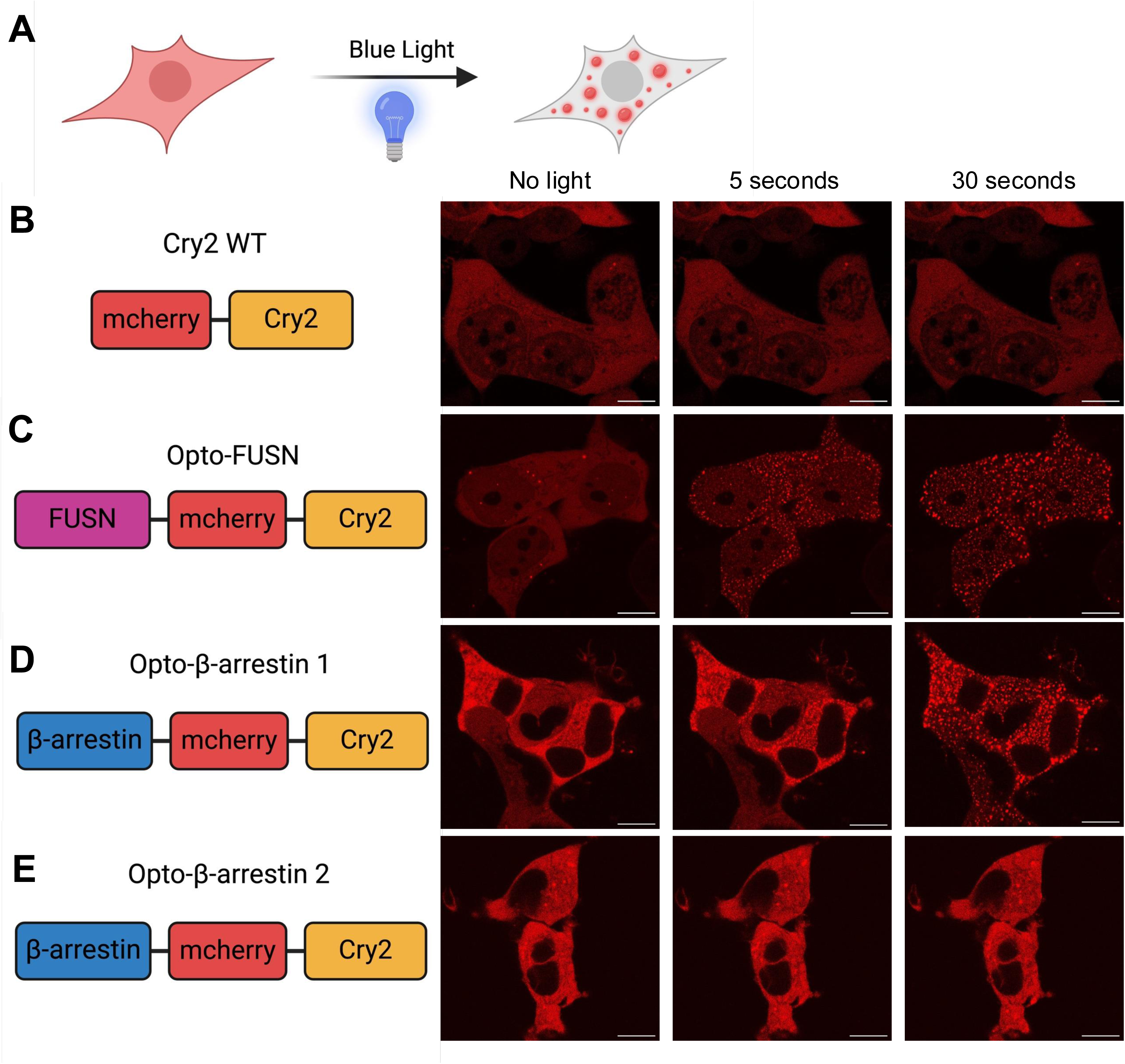
**Development of Opto-**β**-arrestins (A)** Schematic of Optodroplet system. mCherry-Cry2, with either no IDR, FUSN, or β-arrestin 1 or 2, and upon blue light visualization of mCherry. Representative fluorescence of light-activated **(B)** Cry2-WT **(C)** Opto-FUSN **(D)** Opto-β-arrestin 1 **(E)** Opto β-arrestin 2. All cells were transfected with 500ng of respective construct, were shown to have similar expression, and were activated under identical conditions. Scale bars are 10 µm for all images. Confocal microscopy images are representative of n=3.

Notably, the IDRs of β-arrestin are not accessible in the inactive β-arrestin conformation, as the β-arrestin C-terminus is bound to the N-domain^51^. Since IDRs are a major driver of condensate formation, we sought to determine if the inclusion of β-arrestin IDRs alone with Cry2 was also sufficient to drive LLPS. Opto-β-arrestin 1 IDR formed condensates after ∼30 seconds in both the cytosol and the nucleus (**Supplemental Figure 2A and Video S6)**. With the Opto-β-arrestin 2 IDR (**Supplemental Figure 2B and Video S7)**, unlike the full-length β-arrestin 2, we observed condensate formation in both the cytosol and the nucleus. Overall, this data suggests that β-arrestins can undergo LLPS, and the IDRs can drive condensate formation when linked to an oligomerization motif.

### **β**-arrestin oligomerization demonstrates orientation dependence

We next sought to test how different modes of β-arrestin oligomerization could contribute to LLPS. Previous studies have shown that β-arrestins interact through different orientations to generate diverse types of oligomers, such as chains and trimers^8,29,52^. However, it is unclear if GPCRs can promote the orientation-dependent interactions that promote the formation of β-arrestin oligomers. To study this potential process, we used the Nano-BiT system, which relies on the complementation of an 11 amino acid peptide (SmBiT) and a LgBiT fragment^53^. We fused the SmBiT and or LgBiT fragment to either the N-or C-terminus of β-arrestin 2. As there is a low affinity between SmBiT and LgBiT,^54^ any observed signal is driven by the native protein-protein interaction. We transfected different combinations of β-arrestins with these tags on either terminus, e.g. N-N (i.e SmBiT-β-arrestin 2 + LgBiT-β-arrestin 2), N-C (SmBiT-β-arrestin 2 + β-arrestin-2-LgBiT), or C-C (β-arrestin-2-SmBiT + β-arrestin-2-LgBiT). In this assay, we tested the luminescence signal at baseline and the percent change upon agonist stimulation **(Figure 3A)**. Agonist stimulation of the V_2_R resulted in a significant increase in signal with the N-N orientation, followed by N-C and C-C orientations **(Figure 3B)**. At the β_2_AR, agonist stimulation increased the N-C interaction, with a significant decrease in C-C interactions **(Figure 3B).** Both receptors demonstrated a significant decrease in C-C orientation and an increase in N-C and N-N orientations with agonist stimulation. This data suggests different GPCRs promote distinct patterns of β-arrestin oligomerization.

**Figure 3:**
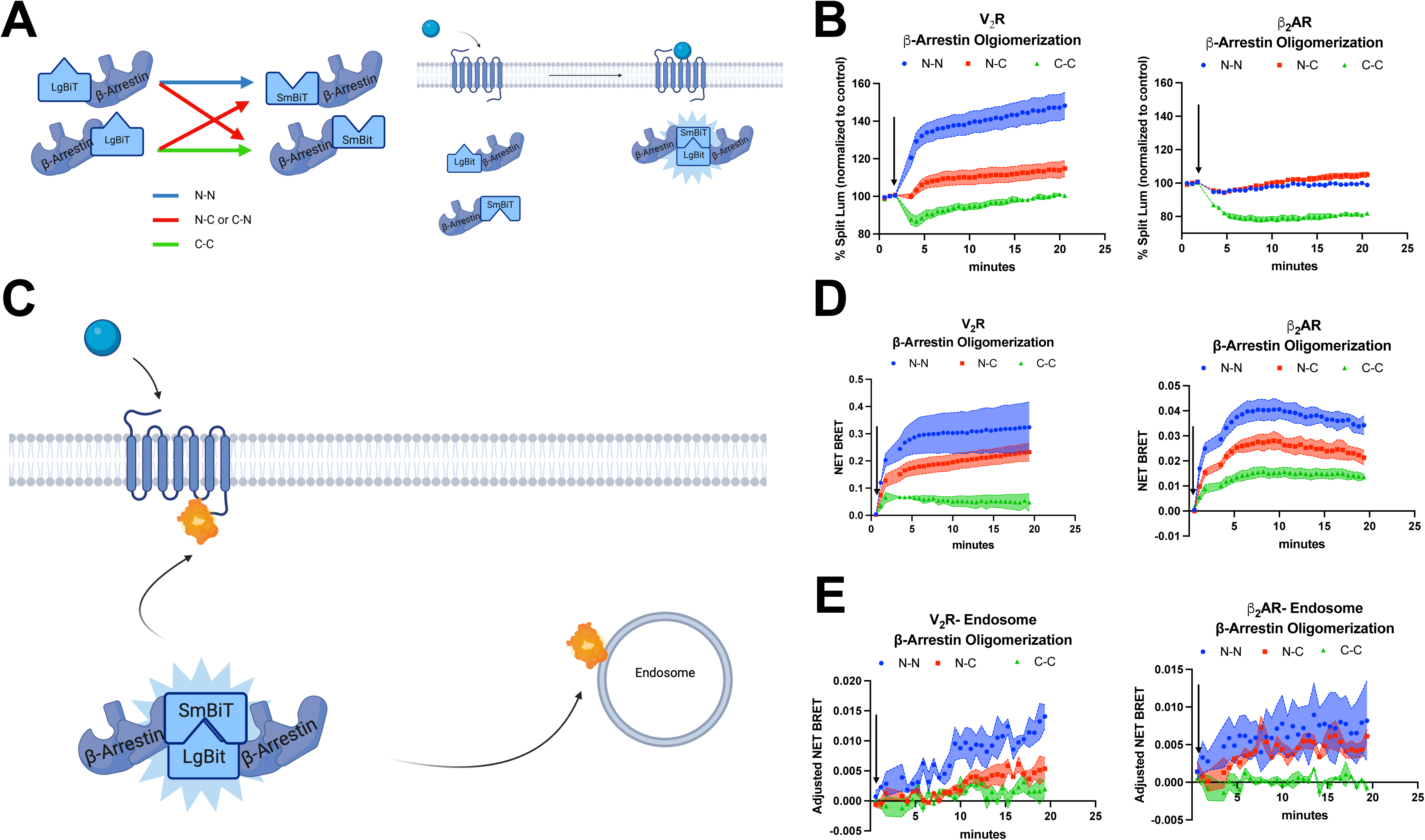
GPCRs regulate. β**-arrestin oligomerization orientation dependence that occurs in proximity to the receptor.** (**A**) β-arrestin split luciferase oligomerization assay. Luciferase fragments (LgBiT or SmBiT) were fused N or C-terminus of β-arrestin 2 to evaluate receptor-induced oligomerization. HEK293 cells were transiently transfected with respective split luciferase β-arrestin, receptors, and stimulated with an agonist or vehicle. (**B**) Receptor-induced β-arrestin-2 oligomerization. Net BRET of cells transfected with V_2_R or β_2_AR and stimulated with AVP (1µM) or isoproterenol (10 µM), respectively. (**C**) β-arrestin-2 oligomerization at receptor or endosome. Cells were either transfected with split luciferase donors (β-arrestin 2) and mKO acceptor (i.e., receptor-or endosome-mKO). (**D**) Net BRET of either V_2_R-mKO or β_2_AR-mKO that were stimulated with AVP (1 µM) or isoproterenol (10 μM), respectively. (**E**) Similar experiment to (**D**) except evaluating β-arrestin 2 oligomerization at endosomes that were stimulated with AVP (1µM) or isoproterenol (10 µM), respectively. For **(B-E)** N=3 separate replicates. The data is shown as mean ± SEM.

These findings suggest that β-arrestin oligomerization was directed in part by GPCR activation, suggesting that it occurs in proximity to the receptor. To assess this, we leveraged an assay for assessing tripartite interactions, NanoBiT bioluminescence resonance energy transfer (BRET),^22,55^ which monitors NanoBiT split-luciferase between two proteins followed by BRET to a fluorescent protein acceptor, monomeric Kusabira orange(mKO). We performed NanoBiT BRET between the SmBiT-and LgBiT-labeled β-arrestins and mKO-tagged receptors (V_2_R-mKO and β_2_AR-mKO) or endosomes (2xFYVE-mKO) **(Figure 3C).** Consistent with our NanoBiT assay, we observed a similar pattern with both the V_2_R-mKO and β_2_AR-mKO, with both receptors displaying an increase in N-N orientation followed by N-C, and C-C β-arrestin orientations. We observed that β_2_AR-mKO promoted nearly ten times lower signal compared to the V_2_R-mKO **(Figure 3D)**. At endosomes, V_2_R promoted predominately N-N, followed by N-C and C-C orientations, while the β_2_AR promoted a similar amount of N-N and N-C oligomerization at endosomes **(Figure 3E)**. We next investigated if β-arrestin can form complexes at other intracellular locations such as the PM (CAAX-mKO) and CCPs (AP2-mKO) **(Supplemental Figure 3A).** For the V_2_R, we found no observable difference in β-arrestin oligomerization at the PM and CCPs (**Supplemental Figure 3B and 3C)**. In contrast, β_2_AR induced a predominately C-C orientation at the PM while it promoted N-C and N-N orientations at CCPs. Overall, these results suggest that receptors can mobilize β-arrestin oligomers to different intracellular compartments.

We next explored if this phenomenon was generalizable to other GPCRs. We selected the Angiotensin II Type 1 Receptor (AT_1_R) and Atypical Chemokine Receptor 3 (ACKR3). Both receptors, in response to agonist stimulation, were able to induce β-arrestin oligomerization with predominately the N-N orientation at the receptor. (**Supplemental Figures 3D and 3E)**. Next, we evaluated if oligomerization had a bystander effect at other receptors (V_2_R and β_2_AR). When the AT1R was stimulated with AngII or ACKR3 was stimulated with WW36, we observed no oligomerization at the non-cognate receptor (V_2_R and β_2_AR, respectively), consistent with oligomerization being promoted at the agonist-activated receptor and not more broadly throughout the plasma membrane (**Supplemental Figures 3D and 3E).** Together, this data demonstrates that numerous GPCRs can promote β-arrestin oligomerization with orientation dependence in proximity to the receptor.

### 1,6 hexanediol diminishes GPCR endocytosis

Phase separation is important for components of the endocytic machinery, such as coating proteins, endocytic adaptors, and membrane lipids^56^ ^57–59^ ^60,61^. Since β-arrestins are critical for GPCR internalization, we hypothesized that internalization is regulated by β-arrestin condensates. We first investigated if 16HD alters the recruitment of β-arrestin to the receptor. Consistent with HEK293 cells pre-treated with 16HD **(Figure 1I),** there was no observable difference between the β-arrestin-GFP vehicle-and 16HD-treated cells. Upon AVP stimulation, β-arrestin-GFP was recruited to the PM, and this recruitment pattern was blunted with pre-treatment with 16HD **(Supplemental Figure 4A)**. In addition, 16HD significantly reduced the initial (∼ 5 minutes) agonist-induced recruitment of both β-arrestin 1 and 2 to different receptors (V_2_R, AT_1_R, and β_2_AR). In contrast, β-arrestin recruitment to the plasma membrane was increased at the V_2_R and AT_1_R, but not the β_2_AR **(Supplemental Figure 4B and 4C).** This difference in recruitment could be attributed to 16HD, since it has peak effectiveness around ∼5 minutes^62^. Overall, our data suggests that 16HD disrupts agonist-induced β-arrestin recruitment.

Since 16HD diminished β-arrestin recruitment, we next tested if GPCR internalization was also inhibited by pre-treatment with 16HD **(Figure 4A-C)**. To control for endogenous β-arrestin expression, we used WT HEK293 cells and studied receptor internalization of two class B GPCRs that are known to require β-arrestins (V_2_R and AT_1_R) **(Figure 4A)**. Incubation with 2.5% or 5% 16HD for 5 minutes resulted in a near-complete ablation of receptor-mediated endocytosis at both the V_2_R and AT_1_R **(Figure 4A).** To confirm these findings, we performed confocal microscopy of HEK293 cells transfected with V_2_R-mKO and AT_1_R-mKO incubated with 2.5% 16HD for 5 minutes and stimulated with their endogenous agonists for 30 minutes. We found receptor-mKO signal was predominately at the plasma membrane, while only vehicle-treated (agonist only with no 16HD) had internalized mKO-labeled receptors (**Figure 4B and 4C)**. These findings are consistent with phase separation being important for GPCR internalization.

**Figure 4:**
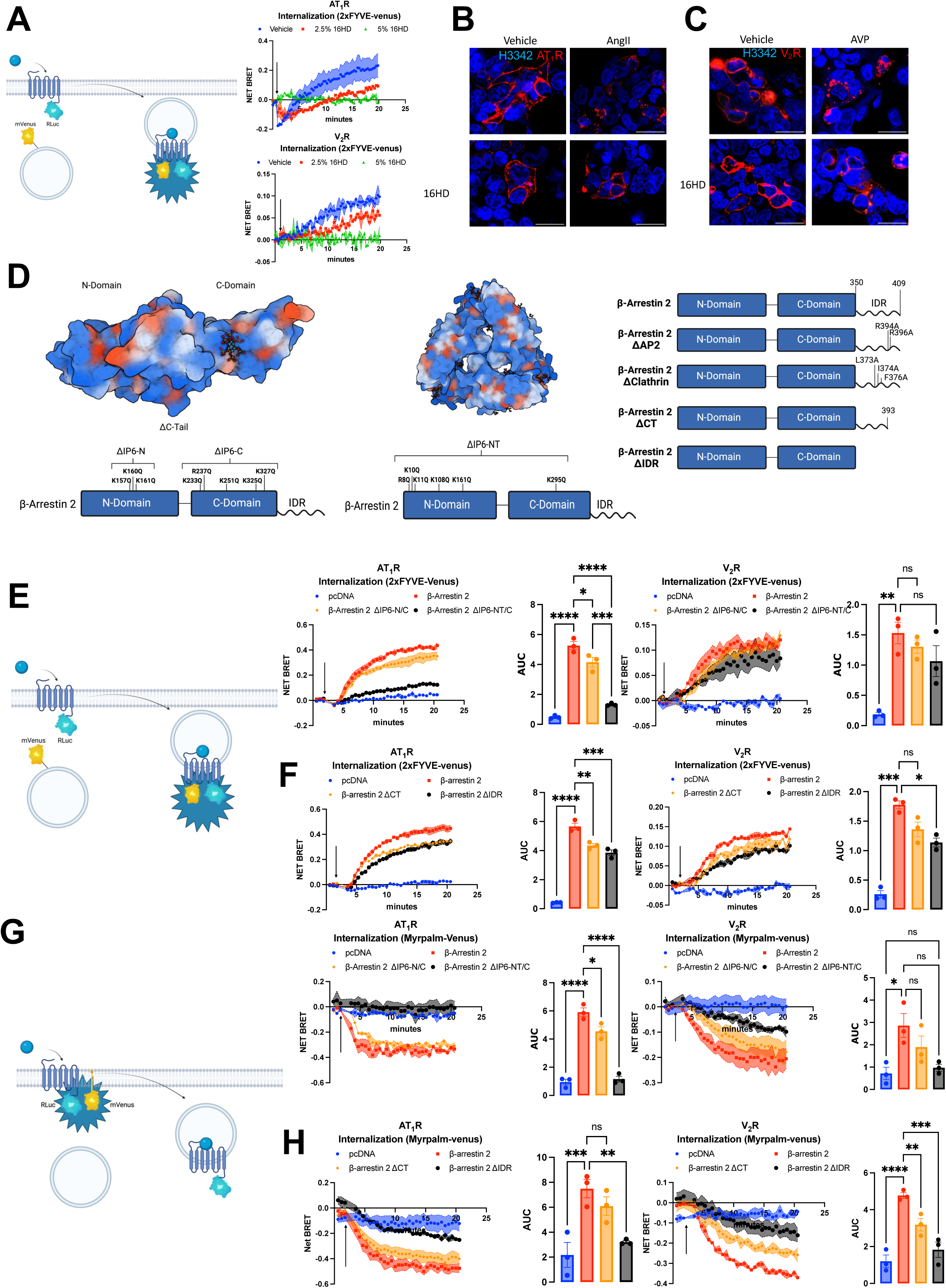
Disruption of. β**-arrestin oligomerization motifs and IDRs impact GPCR internalization.** (**A**) Receptor internalization assay. HEK293 cells were transfected with receptor-Rluc bioluminescent donor, fluorescently-tagged endosome acceptor, and pre-incubated with either vehicle, 2.5% 16HD, or 5% 16HD for 5 minutes. AT_1_R-Rluc was stimulated with AngII (1 μM) and V_2_R-Rluc stimulated with AVP (1uM). (**B-C**) Receptor-mKO transfected cells were treated with vehicle (HBSS) (upper left), stimulated with ligand (upper right), pre-incubated with 2.5% 16HD (lower left), or pre-incubated with 2.5% 16HD and stimulated with ligand (lower right). (**D**) β-arrestin chain, trimeric, and IDR mutants. β-arrestin 1 bound to IP_6_ structure (PDB: 1ZSH). Sequence alignment of β-arrestin-1 and 2 IP_6_ mutagenetic sites: (ΔIP_6_-N, ΔIP_6_-C, and ΔIP_6_-N/C). β-arrestin trimeric structure bound to ΔIP_6_ (PDB: 5TV1) and schematic of the N-terminus trimeric mutant (ΔIP_6_-NT). β-arrestin IDR mutants were designed with MobiDB, β-arrestin 2 IDR extends from 350-418 and includes ΔAP_2,_ ΔClathrin, ΔCT, and ΔIDR. (**E-F**) Receptor internalization assay. Similar to (**A**), except β-arrestin 1/2 KO cells were transfected with β-arrestin mutants AT_1_R: AngII (1 μM) V_2_R: AVP (1 μM). (**G-I**) Receptor internalization assay with Myrpalm-Venus. Similar experiment to **(E-F)** except Venus was fused to the plasma membrane (Mrypalm-Venus). Figures showing the AUC display the mean, SEM, and replicates of the raw kinetic data presented in the preceding panels. *P*LJ≥LJ0.05, \**P =* 0.01–0.05, \*\**P =* 0.001–0.01, \*\*\**P* = 0.0001 to 0.001, \*\*\*\**P*LJ<LJ0.0001 denotes statistically significant differences between AUC by two-way ANOVA. For **(A) (E-I)** N=3 separate replicates. Confocal microscopy images are representative of n=3.

### **β**-arrestin oligomerization and IDR motifs contribute to GPCR internalization

β-arrestins have an architecture with N-and C-domains followed by a C-terminal IDR **(Figure 4D)**. In the N-and C-domains, β-arrestin 1 and 2 have basic residues that bind to IP_6_ and are known to promote oligomer formation in vivo and in vitro ^8,52^ ^28,52^. β-arrestin 1 can form chains^8^ and β-arrestin 2 can form trimers^27^. Since oligomerization can drive LLPS^63,64^, we generated β-arrestin mutants predicted to disrupt IP6 binding sites (ΔIP_6_) that promote oligomerization through chains or trimers. For the chain mutants, we generated three β-arrestin mutants: 1) N-terminus (ΔIP_6_-N); 2) C-terminus (ΔIP_6_-C); and 3) both N and C-terminus (ΔIP_6_-N/C) **(Figure 4D).** Since the ΔIP_6_-C site is the same for the chain and trimer mutants, which share the same IP_6_ site, we generated only the N and C-terminus terminus mutant (ΔIP_6_-NT/C) to disrupt the potential trimer interaction **(Figure 4D).** Therefore, the ΔIP_6_-N/C mutant was referred to as the chain mutant and ΔIP_6_-NT/C as the trimeric mutant. Of note, these mutations did not significantly alter protein expression **(Supplemental Figure 4D)**^65^.

Since IDRs contribute to LLPS, we also sought to investigate the roles of motifs in the β-arrestin-2 IDR. The β-arrestin-2 IDR contains motifs that are important for intracellular localization to CCPs and cytosol^66,67^. We assessed the function of four different β-arrestin 2 IDR mutants, which we named ΔAP2^68^, ΔClathrin^69^, ΔCT^7,70^, and ΔIDR based on the sites that were mutated or deleted **(Figure 4D).** To determine the role of the IP_6_ and IDR β-arrestin mutants in GPCR internalization, we overexpressed these constructs in β-arrestin 1/2 KO HEK293 cells.^71,72^ Internalization was quantified by monitoring BRET between GPCR-RLuc and endosomes tagged with Venus **(Figure 4E).** At the AT_1_R and V_2_R, internalization was not impacted by pcDNA control, but was restored with transfection of WT β-arrestin 2. We found a significant decrease in AT_1_R internalization with the ΔIP_6_-N/C and ΔIP_6_-NT/C mutants compared to WT β-arrestin 2, and a non-significant decrease in V2R internalization with those mutants compared to WT β-arrestin 2. We observed no difference in AT1R or V2R internalization with the ΔIP_6_-N and ΔIP_6_-C mutants compared to WT β-arrestin 2 **(Supplemental Figure 5A).** We next evaluated the contribution of β-arrestin 2 ΔIDR mutants to the internalization of AT_1_R and V_2_R. At the AT_1_R, we observed diminished internalization with all four IDR mutants (ΔAP2, ΔClathrin, ΔCT, and ΔIDR) **(Figure 4F and Supplemental Figure 5B)**. At the V_2_R, three of the four mutants resulted in diminished internalization, except for the ΔCT mutant. We did not observe complete ablation of internalization with ΔIDR mutant, consistent with work showing class B GPCRs utilize motifs in both the IDR and C-lobe of β-arrestins to promote internalization^73^. Next, we employed two different GPCR internalization assays to confirm these results. First, we monitored a loss of BRET signal between the RLuc-labeled receptor and a Myrpalm-labeled Venus (**Figure 4G-H and Supplemental Figure 5C-D)** and performed confocal microscopy **(Supplemental Figure 5E).** Both of these assays exhibited similar results to our endosomal internalization assay. Overall, these results demonstrate that oligomerization motifs and IDRs within β-arrestins contribute to GPCR internalization.

### **β**-arrestin IP_6_ and IDR sites impact GPCR-mediated signaling

We next tested the consequences of oligomerization motifs and IDRs on signaling downstream of GPCRs. To do this, we tested the impact of β-arrestin oligomerization and IDR mutants on activity of extracellular signal-regulated kinase 1 and 2 (ERK1/2) kinases, which are key regulators of cell proliferation, survival, and apoptotic signaling^74^. Since AT_1_R is an inducer of ERK activity,^3^ we sought to evaluate ERK1/2 activity in different intracellular locations. We employed a BRET-based ERK activity reporter (EKAR) targeted to different locations^7576^ **(Figure 5A-B)**. In HEK293 cells, titration of 16HD resulted in a decrease in AngII-induced cytosolic and nuclear ERK activity **(Figure 5B)**, which was confirmed by Western blot **(Supplemental Figure 6A-B).** While this data is consistent with LLPS regulating basal and GPCR-mediated ERK signaling in different intracellular compartments, other potential effects of 16HD cannot be excluded in these studies^77^.

**Figure 5:**
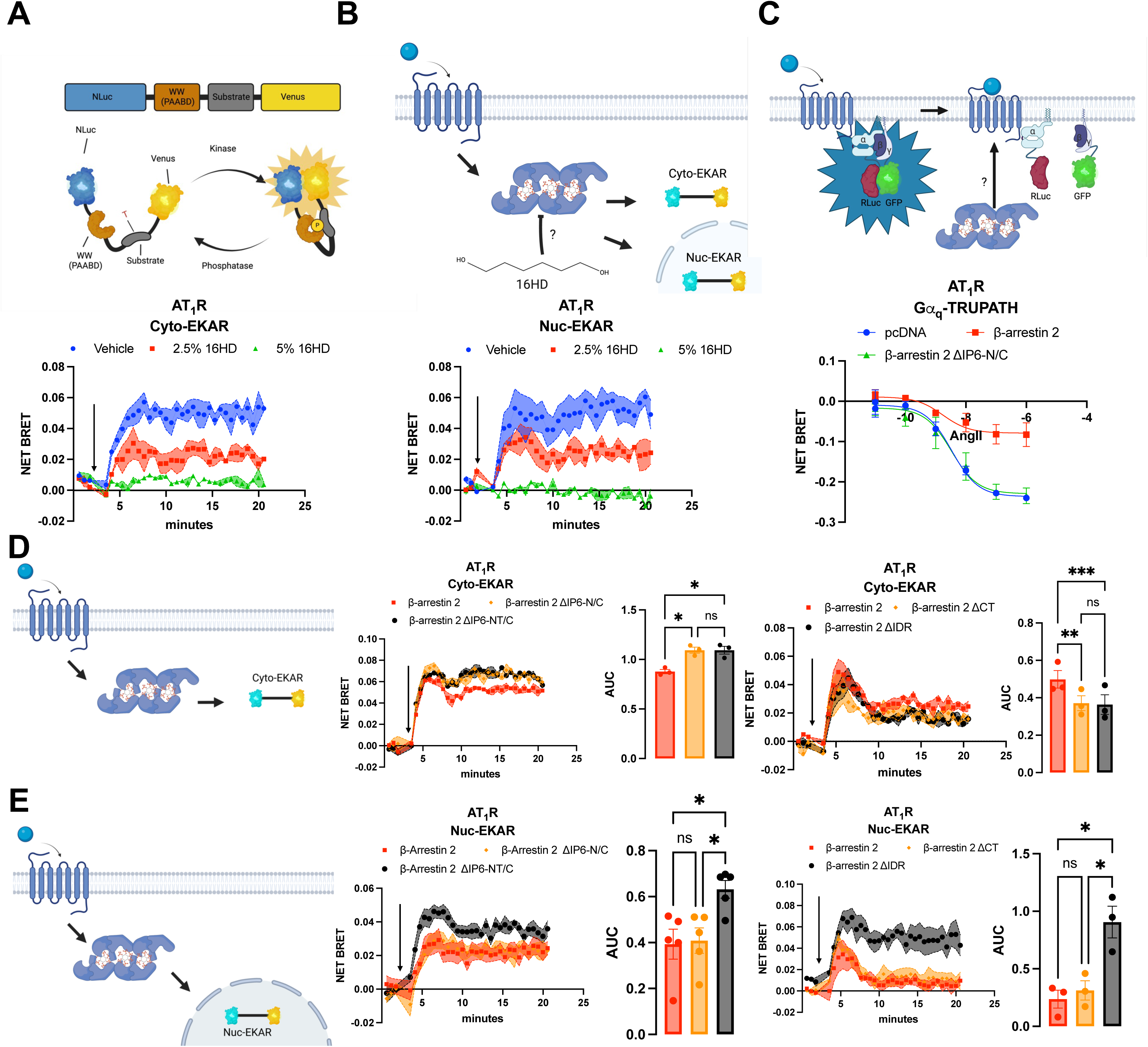
β**-arrestin IP_6_ and IDR mutants influence GPCR signaling. (A)** Schematic of BRET-based ERK biosensor that utilizes ERK-specific substrate that upon phosphorylation induces a conformational change and an increase in BRET. (**B**) β-arrestin-mediated cytoplasmic and nuclear ERK sensor readout in cells incubated with 16HD. Net BRET readout from localized ERK biosensor in HEK293 cells that were transfected with AT_1_R, stimulated with AngII (1 μM) or vehicle, pre-treated with 16HD for 5 minutes. (**C**) GIZI_q_ TRUPATH. HEK293 cells were transfected with AT_1_R: GIZI_q_ TRUPATH components: β-arrestin WT and mutants and stimulated with a AngII dose-response. (**D**) Kinetic data and AUC quantification of cytoplasmic ERK sensor with β-arrestin WT and mutants after stimulation with AngII (1 μM) (**E**) Similar to experiment (**D**) except nuclear ERK monitored with Nuc-EKAR. Figures showing the AUC display the mean, SEM, and replicates of the raw kinetic data presented in the preceding panels. *P*LJ≥LJ0.05, \**P =* 0.01–0.05, \*\**P =* 0.001–0.01, \*\*\**P* = 0.0001 to 0.001, \*\*\*\**P*LJ<LJ0.0001 denotes statistically significant differences between AUC by two-way ANOVA. For **(A-E**), N=3, except Nuc-EKAR (β-arrestin ΔIP_6)_) N=5 replicates per condition Cyto, cytoplasmic. Nuc, Nuclear. Data shown as mean ± SEM.

One of the chief functions of β-arrestin is to desensitize the receptor by sterically inhibiting G-proteins from interacting with the receptor^78^. Since β-arrestin ΔIP_6_ oligomeric mutants exhibited different internalization patterns, we next sought to investigate how this influenced G-protein activation^79,80^ **(Figure 5C and Supplemental Figure 6C).** In HEK293 cells, GIZI_q_ dissociation for AT_1_R was significantly diminished by the expression of WT β-arrestin 1 or 2. Interestingly, the ΔIP_6_-N/C mutant had no significant effect on G protein signaling compared to pcDNA control **(Figure 5C and Supplemental Figure 6C)**. We then sought to investigate if the IP_6_ and IDR motifs in β-arrestins regulate AT_1_R-mediated ERK signaling^81^ **(Figure 5D-E and Supplemental Figure 6D-E)**. Consistent with previous studies^4^, overexpression of β-arrestin 2 in HEK293 cells resulted in less cytoplasmic ERK compared to pcDNA control **(Supplemental Figure 6D-5E)**. There was no decrease in cytoplasmic ERK activity with the majority of β-arrestin ΔIP_6_ mutants except for β-arrestin ΔIP_6_-C. In contrast, all four β-arrestin 2 IDR mutants (ΔCT, ΔIDR ΔAP2, ΔClathrin) resulted in a small reduction in cytoplasmic ERK **(Figure 5D and Supplemental Figure 6D).** In the nucleus, transfection of β-arrestin 2 ΔIP_6_-NT/C resulted in more ERK activity compared to WT β-arrestin 2 **(Figure 5E and Supplemental Figure 6E).**

Most of the IDR mutants resulted in a decrease in nuclear ERK activity, except for the ΔIDR mutant, which increased activity **(Figure 5E and Supplemental Figure 6E)**. These findings suggest contrasting roles for specific IP6 binding and IDR motifs in the regulation of GPCR signaling, some of which may be related to their regulation of LLPS and others which may be related to other β-arrestin-mediated functions.

### Structure of a 2:2 ADGRE1:**β**-arrestin 1 complex

To provide insights into the structural mechanisms that underlie β-arrestin oligomerization, we determined the structure of a GPCR that constitutively interacts with β-arrestins. Previous work has demonstrated that the adhesion receptor (aGPCR) ADGRE1 displays high basal recruitment of β-arrestin^82^. We found that ADGRE1 formed a stable complex with β-arrestin1 (**Supplemental Figure 7A-C**). To facilitate ADGRE1-β-arrestin 1 complex formation, wild-type full-length mouse ADGRE1 (residues 1-386) with its intact C-terminal tail was expressed with a constitutively active truncated cysteine-free bovine β-arrestin 1 (residues 1-376). An engineered scFv30 was fused to the C-terminal of β-arrestin 1 to ensure efficient expression and stabilization of the ADGRE1-β-arrestin 1 complex and the receptor was phosphorylated in vitro by GRK2. For structure determination, ADGRE1, β-arrestin 1, and GRK2 were co-expressed in Sf9 cells using the Bac-to-Bac system. The complex was purified and cryo-EM data were collected on a 300 kV Titan Krios G4 microscope equipped with a Selectris X imaging filter and Falcon 4 detector. A total of 23,906 movies were collected with a final reconstruction of a global map at 3.58 Å resolution (**Supplemental Figure 7D-H**).

The cryo-EM structure of the ADGRE1-β-arrestin 1 complex reveals that ADGRE1 forms a parallel dimer, with two ADGRE1 symmetrically arranged protomers engaging β-arrestin 1 (**Figure 6A** and **B**). The resolution was sufficient to resolve most regions of the complex (**Supplemental Figure 7I**), although the extracellular N-terminal fragment (NTF) was not visible, likely due to its intrinsic flexibility, leading to density averaging. The dimer interface is primarily mediated by transmembrane helix 1 (TM1), transmembrane helix 2 (TM2), and helix 8 (H8), forming extensive intermolecular contacts that stabilize the dimeric assembly (**Fig. 6C**). This parallel dimerization mode differs from previously characterized GPCR-arrestin structures, which are predominantly monomeric. Paralleling to our work, a recent study of mGlu-arrestin structure also exhibits a dimerization mode^83^, suggesting a potential distinct mechanism for GPCR signaling.

**Figure 6:**
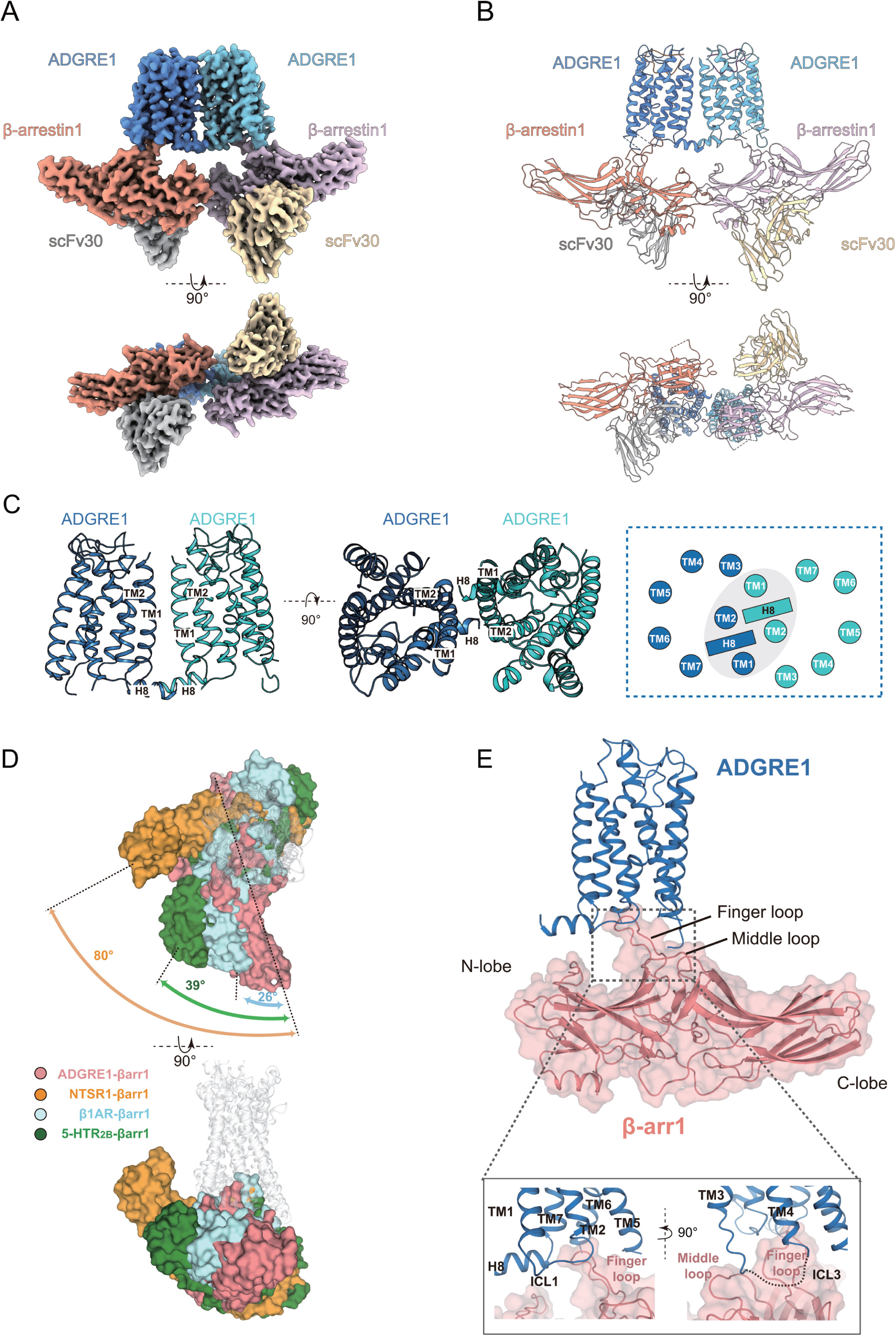
**Structure of the ADGRE1-**β**arrestin 1 complex reveals a dimeric organization and potential for higher-order organization.** (**A, B** Overview cryo-EM map (A) and ribbon representation (B) of ADGRE1-βarrestin 1 complex. Note that ADGRE1 forms a parallel dimer, with two ADGRE1 symmetrically arranged protomers engaging β-arrestin1. (**C**) Intermolecular contacts that stabilize the dimeric ADGRE1 assembly are highlighted, with a schematic cartoon illustrating the close proximity between TM1, TM2, and H8 shown within the dashed box. (**D**) β-arrestin 1 is positioned nearly perpendicular to the receptor’s central axis and parallels to membrane, forming a nearly 90° angle relative to the membrane, in contrast to other GPCR:β-arrestin 1 structures. (**E**) β-arrestin 1 interacts with ADGRE1 through three primary interfaces: the 7TM core, ICL1, and ICL2.

The two ADGRE1-β-Arrestin1-scFv30 complexes in the structure are nearly identical, with a root-mean-square deviation (RMSD) of 0.44 Å for the ADGRE1s and 0.42 Å for the β-arrestins, respectively. One complex exhibited slightly better-defined density and was therefore used as the model for detailed analysis of receptor-arrestin interactions. Both ADGRE1 and β-arrestin 1 adopt active conformational states in the ADGRE1-β-arrestin 1 complex. Compared with the AlphaFold-predicted inactive structure of ADGRE1, the seven-transmembrane (7TM) domain of ADGRE1 in the β-arrestin 1-bound state adopts an active conformation, characterized by a pronounced outward displacement of TM6, a hallmark of GPCR activation (**Supplemental Figure 8A-B**). Similarly, the activation of β-arrestin 1 is marked by a 15° rotation of the C-lobe relative to the N-lobe, as well as the adoption of an active conformation in the central crest loops, distinguishing it from its inactive form (**Supplemental Figure 8C-8D**).

### The GPCR:**β**-arrestin 1 engagement mode allows for **β**-arrestin oligomerization

The ADGRE1-β-arrestin 1 complex adopts an engagement mode distinct from previously characterized GPCR-β-arrestin 1 structures, including NTSR1-β-arrestin 1, β_1_AR-β-arrestin 1, 5HT2B-β-arrestin 1, CB1-β-arrestin 1, V_2_R-β-arrestin 1, and M_2_R-β-arrestin 1 (**Fig. 6D**, and **Supplemental Figure 8E**). A key distinction lies in the orientation of β-arrestin 1 relative to the receptor and the membrane. Structural alignment of ADGRE1-β-arrestin 1 complex and previously reported GPCR-β-arrestin 1 complexes revealed that β-arrestin 1 exhibits a significantly different orientation in the ADGRE1 complex. Measurement of the relative rotation of β-arrestin 1 revealed an engagement angle variation ranging from 21° to 80° compared to its positioning in other GPCR-β-arrestin 1 complexes (**Figure 6D** and **Supplemental Figure 8E**). In addition, in contrast to previously reported GPCR-β-arrestin 1 complexes, where the C-edge of β-arrestin 1 typically interacts with the membrane due to an upward tilt of its C-terminal region, β-arrestin 1 in the ADGRE1 complex adopts a distinct orientation (**Fig. 6E** and **Supplemental Figure 8F**). Rather than tilting toward the membrane, β-arrestin 1 is positioned nearly perpendicular to the receptor’s central axis and parallels to membrane, forming a nearly 90° angle relative to the membrane. This non-canonical binding pose distinguishes ADGRE1 from other GPCR-arrestin complexes The ADGRE1-β-arrestin 1 complex reveals distinct interaction interfaces involving the transmembrane (TM) core and intracellular loops (ICL1 and ICL2) (**Figure 6E**). Unlike previously reported GPCR-arrestin complexes, where the phosphorylated receptor C-tail typically binds to the positively charged N-lobe groove of arrestin, this structure reveals an alternative mode of interaction: the phosphorylated C-tail of ADGRE1 is absent, and instead, β-arrestin 1 engages the receptor’s intracellular loops and transmembrane bundle. In this structure, β-arrestin 1 interacts with ADGRE1 through three primary interfaces the 7TM core, ICL1, and ICL2 (**Figure 6E**). The finger loop of β-arrestin 1 inserts into the receptor’s 7TM bundle, interacting with a pocket formed by ICL1, TM2, TM3, ICL2, TM6, and TM7 (**Figure 6E**).

Specific hydrophobic and polar interactions are observed between β-arrestin 1 finger loop residues R65, E66, D67, D69, V70 and ADGRE1 residues N677^ICL1^, H678^ICL1^, H683^2.50^, M738^3.57^, S749^ICL2^, L841^6.38^, K844^6.41^, H889^7.57^ located in the cytoplasmic regions of TM2, TM3, TM6, TM7, ICL1 and ICL2 (**Supplemental Figure 7J-K**). Additionally, while part of ICL2 is not fully resolved in the density map, it likely interacts with the middle loop of β-arrestin 1 through hydrophobic and polar contacts (**Figure 6G**-**H**). Consistent with these observations, mutations in key residues, including N677^2.44^, H683^2.50^, H889^7.57^ to Ala, significantly reduced β-arrestin 1 recruitment, further supporting the structural findings (**Supplemental Figure 7L**).

### ADGRE1 promotes **β**-arrestin condensate formation

While the β-arrestins in the ADGRE1:β-arrestin 1 structure are not directly interacting, they are in close proximity to one another with the N-terminus of one β-arrestin in close proximity (∼8 Å) to the N-terminus of the other β-arrestin (**Figure 6A-6B**). To determine if this orientation could promote β-arrestin interaction, we performed Metadynamics (MD) simulations of β-arrestin 1. We found that the conformational state of N-N orientation of the ADGRE1-β-arrestin-1 dimer observed in the cryoEM structure could be captured by our Metadynamics simulations (**Figure 7A and Supplemental Figure 9A-9B**). Our simulation results further suggested potential direct N-N orientation dimerization interface of the ADGRE1:β-arrestin-1 (**Figure 7B-7E**). Notably, polar or hydrogen bonds were present between side chains of interface residues in the N-lobe of arrestins. The key residue interaction pairs involved including K10-E46, K11-N15, N15-R161, K17-T19, Y21-K17, D44-K10/K11, Y47-K160, K49-E367, R51-E369, and E110-K107. In total, 10 residues of one protomer engaged with 11 residues of the other promoter of β-arrestin 1. Although the two arrestin protomers in the ADGRE1-β-arrestin 1 dimer conformation do not form permanent interactions, the above-listed interactions were supported by the frequency of contacts formed between these interface residues (**Supplemental Figure 9C**). This supports a model in which the observed ADGRE1:β-arrestin-1 conformation can support the formation of β-arrestin oligomers.

**Figure 7:**
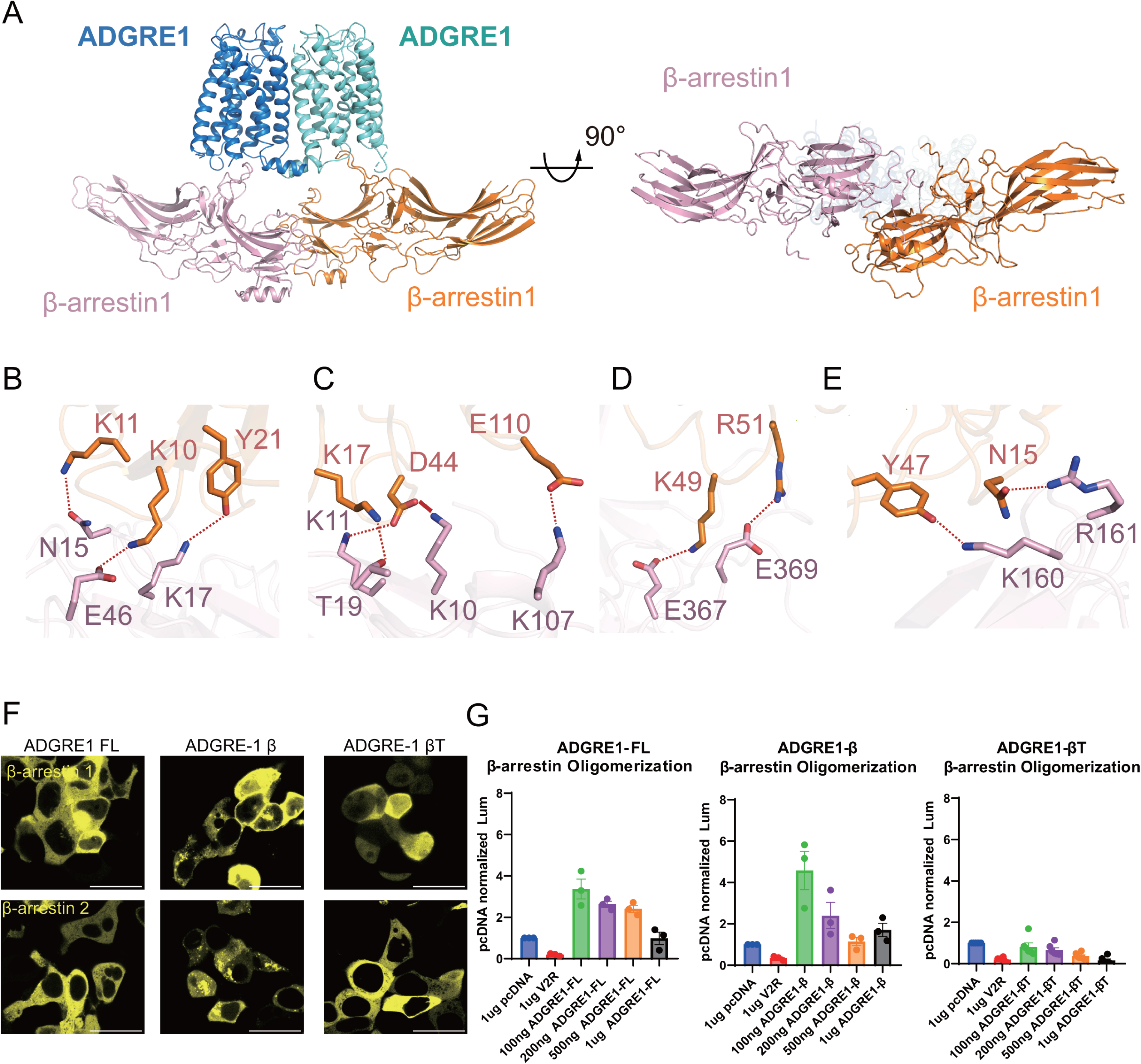
**Metadynamics simulations of ADGRE1–**β**-arrestin 1 dimerization. (A)** Structural representation of the ADGRE1–β-arrestin 1 dimer obtained from Metadynamics simulations. The N-N orientation led to direct β-arrestin interactions. **(B–E)** Detailed interactions at the dimerization interface, highlighting key amino acid residues involved in inter-protein contacts. Red lines represent hydrogen bonds or polar interactions **(F)** HEK293 cells were transfected with either β-arrestin 1 or β-arrestin 2-YFP and respective ADGRE receptor **(G)** β-arrestin split luciferase oligomerization assay. Luciferase fragments were fused N-terminus of β-arrestin 2 to evaluate basal N-N oligomerization. For **(G)** N=3 separate replicates. The data is shown as mean ± SEM. Scale bars are 25 µm for all images. Confocal microscopy images are representative of n=3.

We then tested if ADGRE1 could promote β-arrestin condensate formation when the full length (ADGRE1-FL), constitutively active (ADGRE1-β) or Stächel peptide-deleted (ADGRE1-βT) receptors **(Figure 7F)** were overexpressed in HEK293T cells. With overexpression of ADGRE1-FL, we observed a diffuse cytosolic pattern of β-arrestin expression, consistent with a receptor with low constitutive activity **(Figure 7F)**. In contrast, overexpression of the constitutively active receptor ADGRE1-β promoted the formation of condensates **(Figure 7F).** Lastly, removal of the Stächel peptide, resulting in a “dead” receptor, eliminated puncta formation **(Figure 7F)**. Next, we sought to investigate if ADGRE1 promoted the N-N orientation of β-arrestin oligomers using a NanoBiT β-arrestin oligomerization assay. Titration of ADGRE1-FL and ADGRE1-β resulted in nearly a three and five-fold increase in oligomerization, whereas ADGRE1-βT did not change oligomerization **(Figure 7G)**. Together, these support a model in which active ADGRE1 promotes the formation of β-arrestin oligomers in an N-N orientation.

## DISCUSSION

Our studies support a model where β-arrestins form condensates at baseline and after GPCR stimulation to regulate GPCR function. The formation of condensates that compartmentalize GPCR signaling potentially explains how β-arrestins are capable of interacting with hundreds of effector proteins to regulate GPCR signaling. We provided multiple lines of evidence to support this model. We demonstrated that β-arrestins form condensates with split fluorescent proteins when activated by GPCR activation and by using optogenetic systems. We then showed the impact of specific oligomerization motifs and IDRs on β-arrestin activity at GPCRs, including internalization and signaling. Lastly, we provide structural evidence from an ADGRE1:β-arrestin 1 cryoEM structure that β-arrestins can engage with receptors in a mode that promotes β-arrestin oligomerization, which we support with MD simulations and assessment of β-arrestin condensate formation and oligomerization. Leveraging these orthogonal approaches, our results support a new paradigm for how β-arrestin condensates can for condensates that contribute to GPCR function.

In recent years, biomolecular condensates have been uncovered as playing important roles in many aspects of intracellular signaling. Receptors themselves can form condensates such as RTKs and T-cell receptors.^36,37^ In addition, different signaling effectors such as PKA and WNK have been shown to undergo LLPS^35,38^. Macromolecules enriched in condensates exhibit intra-and intermolecular multivalent interactions that are mediated by motifs that promote oligomerization and the presence of IDRs,^32^ features that are present in β-arrestins. The observed differences between β-arrestin 1 and 2 in condensate formation in our assays could be related to the ability of β-arrestin 1 to form “infinite chains”^8^ while β-arrestin 2 forms trimers^27^. Interestingly, the IDR of either β-arrestin was sufficient to drive LLPS formation in the optogenetic system. β-arrestins with a functional IDR resulted in increased cytosolic ERK signaling, and the monomeric form (truncated IDR and ablation of IP_6_ sites) of β-arrestin resulted in increased nuclear signaling. This is consistent with previous studies that have proposed that β-arrestin 2 may sequester ERK in the cytosol^4,81^. β-arrestins 1 and 2 are known to interact with hundreds of proteins important for signaling, transcription, and cytoskeleton^5^.

Future studies will characterize how β-arrestin condensates and how different components such as oligomerization and IDR contribute to regulate GPCR signaling.

We also provide the first structure of an aGPCR bound to a β-arrestin, which demonstrated an aGPCR dimer, with each monomer interacting with β-arrestin 1 in a unique mode compared to previous structures. The β-arrestins were in close proximity to each other, and using MD simulations, the observed conformation of the β-arrestins could support the formation of an N-N interaction. This was supported by the ability of a constitutively active ADGRE1-β to promote condensates and N-N oligomers. Some of the first evidence that β-arrestin could hetero-and homo-oligomerize was in cells and displayed an orientation dependence bias at baseline^29^. Except for arrestin 4 (visual arrestin), all arrestins have been shown to oligomerize, but the oligomers they form when purified are different^28^. Notably, β-arrestin 1 and 2 mutants that are unable to bind IP_6_ do not appear to oligomerize in vivo^30^.

Recent structural work has shown phosphopeptide C-tails (V_2_Rpp, C5a1Rpp, and M_2_Rpp) induce a trimeric β-arrestin structure^84,85^, suggesting oligomerized β-arrestin could interact with GPCR C-tails. We show that β-arrestin oligomerization occurs near receptors that display a common orientation (N-N > N-C > C-C orientations) and that oligomerization mutants reduced internalization. Further studies are required to assess the functional significance of these different orientations.

In summary, our findings demonstrate that β-arrestin can form condensates that regulate GPCR function. This finding addresses open questions regarding the function of β-arrestin oligomerization on GPCR function.^28,86,87^ Given the importance of β-arrestins in GPCR regulation and drug discovery, these findings have translational and clinical implications, as context-dependent targeting of these condensates could be used to regulate physiologically relevant GPCR signaling^2^ that may aid in the development of more efficacious GPCR therapeutics.

### Limitations of the Study

To pharmacologically investigate LLPS formation, we used 16HD, a non-specific inhibitor of hydrophobic regions, which has been shown to non-specific effect on kinase function^77^.

Oligomerization can be difficult to quantify, so for many of our studies, we leveraged orthogonal split assays (i.e., split-GFP^88^ and split luciferase^54^) in HEK293 cells^89^. Many experiments relied on the overexpression of components that have been genetically modified or have the insertion of fluorescent or luciferases. To investigate the role of IP_6_ on β-arrestin oligomerization, we mutated known IP_6_ binding sites in β-arrestins, as modulating the intracellular concentration of IP_6_ can be challenging. The expression of β-arrestin IP_6_ mutants were normalized to endogenous HEK293 cell expression. It is feasible that IP_6_ sites may have multiple functions, which were also disrupted with mutation.

## Acknowledgments

We thank Robert J. Lefkowitz, Sudha Shenoy, Laura Wingler, Claudia Lee, Uyen Pham and Dylan Eiger and for their helpful discussion and thoughtful feedback. The cryo-EM data were collected at the Biomedical Research Center for Structural Analysis, Shandong University. We thank the Translational Medicine Sharing Platform of the Advanced Medical Research Institute, Shandong University, for technical support. The authors gratefully acknowledge the Duke Light Microscopy Core Facility for their support and assistance in this work. Graphical Figures were made using BioRender. This work was supported by an American Heart Association predoctoral fellowship 23PRE1011306(P.J.A), 23PRE1019796 (U.P), Duke Medical Scientist Training Program (PJA), R01GM122798 (S.R.), K08HL114643 (S.R.), and a Mandel Seed Award (S.R.).

## Author Contributions

Conceptualization, P.J.A, P.X., J.P S, S.R.

Methodology, P.J.A, P.X, A.K, J.A,Y.Z, T.Z, C.K, K.Y, L.Q, W.D, S.L, B.P, A.K,O.S, C.J J. S,

Investigation, P.J.A, P.X, A.K, J.A,Y.Z, T.Z, C.K, K.Y, L.Q, W.D, S.L, B.P, A.K,O.S, C.J J. S

Writing-Original Draft, P.J.A., P.X., A.K, J.P S.R

Writing-Reviewing and Editing, P.J.A, P.X, A.K, J.A,Y.Z, T.Z, S.L, JPS

Visualization, P.J.A, S.R.

Supervision and Funding Acquisition, P.J.A, J.P, S.R

## Declaration of Interests

None

## Supplemental Figures

**Supplemental Figure 1:** β**-arrestin-1 and 2 form condensates. (A)** Expression of GFP_11_ or sfGFP_1-10_ individual components in HEK293 cells **(B)** Titration of β-arrestin 1 and 2 eGFP. HEK293 cells were transfected with either 100, 200, or 400ng of β-arrestin 2 eGFP **(C)** Similar to **(B),** but transfection of β-arrestin split-GFP constructs **(D)** Immunoblot of β-arrestin eGFP and β-arrestin split-GFP titrations. Scale bars are 25µm for all images. Confocal microscopy images are representative of n=3.

**Supplemental Figure 2: Characterization of Opto** β**-arrestin IDR.** The IDR of β-arrestin 1 or 2 was fused to the N-terminus of Cry2-mCherry. Representative fluorescence images of light-activated **(A)** Opto β-arrestin 1 IDR (**B)** Opto β-arrestin 2 IDR. IDR sites were identified from MobiDB. All cells shown have similar expression of constructs and are activated under identical conditions. Scale bars are 10 µm for all images. Confocal microscopy images are representative of n=3.

Supplemental Figure 3: **β**-arrestin oligomerization occurs in proximity to the receptor.

**(A)** β-arrestin-2 subcellular oligomerization assay. Luciferase fragments (LgBiT or SmBiT) were fused to the N or C-terminus of β-arrestin 2 to evaluate receptor-induced oligomerization **(B)** BRET of β-arrestin 2 oligomerization was evaluated at the plasma membrane (CAAX-mKO) upon agonist stimulation of AVP (1µM) or Isoproterenol (10 µM). **(C)** Similar experimental design to **(B),** except using AP2-mKO to label clathrin-coated pits upon agonist stimulation.

**(D)** β-arrestin oligomerization at AT_1_R-mKO or V_2_R-mKO both treated with AngII (1µM). **(E)** ACKR3-mKO or V_2_R-mKO mediated β-arrestin-2 oligomerization treated with WW36 (1µM). For **(B-E)** N=3 separate replicates. The data shown are the mean ± SEM

**Supplemental Figure 4: Characterization of 16HD on** β**-arrestin recruitment. (A)** β-arrestin recruitment to the receptor. HEK293 cells were transfected with β-arrestin-GFP, untagged V_2_R, and were treated with either vehicle or 2.5% 16HD, vehicle or agonist AVP (1μM) **(B)** β-arrestin split luciferase recruitment to V_2_R-LgBiT stimulated with AVP, (1 µM), β _2_AR-LgBiT with Isoproterenol (10 µM), AT_1_R-LgBiT with AngII (1 µM) **(C)** β-arrestin split luciferase recruitment to the plasma membrane. A similar experiment to **(B)** except the LgBiT was localized to the plasma membrane (CAAX-LgBiT). **(D)** Immunoblot and quantification of HEK293 cells transfected with β-arrestin 2 and ΔIP_6_ mutants. For **(A-D)** N=3 replicates per condition **(A)** images are representative of n=3.

S**upplemental Figure 5: Additional characterization of the effects of** β**-arrestin mutants on GPCR internalization. A-B)** Endosome receptor internalization assay of the V_2_R stimulated with AVP (1 µM) and AT_1_R with AngII (1 µM) **C-D)** Receptor internalization assay, similar experiment to **(A-B),** except the fluorescent acceptor was localized to the plasma membrane (Myrpalm-venus) **(E)** β-arresin ½ KO cells were transfected with Receptor-mKO and either pcDNA3.1 or β-arrestin ΔIP_6_ mutants treated with saturating amount of ligand for 20 minutes. **A-D)** Figures showing the AUC display the mean, SEM, and replicates of the raw kinetic data presented in the preceding panels. *P*LJ≥LJ0.05, \**P =* 0.01–0.05, \*\**P =* 0.001–0.01, \*\*\**P* = 0.0001 to 0.001, \*\*\*\**P*LJ<LJ0.0001 denotes statistically significant differences between AUC by two-way ANOVA / For **(A-D)** N=3 replicates per condition **(E)** images are representative of n=3.

**Supplemental Figure 6: Additional characterization of** β**-arrestin mutants on GPCR signaling (A)**Localization of β-arrestin 2 split-GFP and ERK-mKO. HEK293 cells were transfected with AT_1_R, β-arrestin-2 split-GFP system, and stimulated with AngII (1μM).(**B)** Oligomeric β-arrestin-2 interaction with ERK. Corrected Net BRET of β-arrestin/β-arrestin: ERK-mKO upon incubation of either 2.5 or 5% 16HD. **(A)** ERK Immunoblots from HEK293 cells incubated with 16HD or vehicle. **(B)** ERK Immunoblots from HEK293 cells that were transfected with AT_1_R, stimulated with AngII (1uM), and incubated for 5 minutes of 16HD or vehicle **(C)** GIZI_q_ TRUPATH. HEK293 cells were transfected with AT_1_R: TRUPATH components: β-arrestin WT and mutants and stimulated with a AngII dose-response **(D)** β-arrestin mediated cytoplasmic ERK sensor. Cyto-EKAR BRET readout from cells that were transfected with AT_1_R, β-arrestin ΔIP_6_ or IDR mutants and stimulated with AngII (1uM) **(E)** Similar to panel **(D)** but Nuc-EKAR. Figures showing the AUC display the mean, SEM, and replicates of the raw kinetic data presented in the preceding panels. *P*LJ≥LJ0.05, \**P =* 0.01–0.05, \*\**P =* 0.001–0.01, \*\*\**P* = 0.0001 to 0.001, \*\*\*\**P*LJ<LJ0.0001 denotes statistically significant differences between AUC by two-way ANOVA. For (**A-E),** N=3, except Nuc-EKAR (β-arrestin ΔIP_6)_) N=4. Data shown are the mean ± SEM

**Supplemental Figure 7: Workflow for structure determination of ADGRE1-**β**-arrestin1 complex. (A)** Snake representation of ADGRE1 construct used in this study. **(B-C)** Representative SDS-PAGE analysis of in vitro reconstituted ADGRE1-β-arrestin1 complex (B) and corresponding elution profiles from a Superose 6 Increase 10/300 column, note that the red arrow indicats the peak fractions that were collected and subsequently used for data collection

(C). **(D-E)** Representative cryo-EM micrograph (D) and two-dimensional (2D) class averages (E) of ADGRE1-β-arrestin1 particles. **(F)** Flowchart of cryo-EM data processing of ADGRE1-β-arrestin1 complex. **(G)** 3D density map colored according to local resolution (Å) of the ADGRE1-β-arrestin1 complex. **(H)** Fourier shell correlation curves for the final 3D density map of ADGRE1-β-arrestin1 particles. At the FSC 0.143 cut-off, the overall resolution for the map is 3.58 Å. **(I)** Cryo-EM density of transmembrane helices of ADGRE1, and figure loop, middle loop, and lariat loop of β-arrestin1 determined in ADGRE1-β-arrestin1 cryo-EM structure.

Hydrophobic and polar interactions are observed between (**J**) β-arrestin 1 finger loop residues R65, E66, D67 with ADGRE1 residues N677^ICL1^, H678^ICL1^, and S749^ICL2^; and (**K**) β-arrestin 1 finger loop residues D69 and V70 with ADGRE1 residues H683^2.50^, L841^6.38^, K844^6.41^, and H889^7.57^. (**L**) Mutations in key residues N677^2.44^, H683^2.50^, H889^7.57^ to Ala, significantly reduced β-arrestin 1 recruitment compared to wild type.

**Supplemental Figure 8: Structural analysis of the ADGRE1** β**-arrestin1 complex. (A-B)** Structural comparison of the transmembrane domain (TMD) between β-arrestin1-coupled ADGRE1 (blue) and the AlphaFold-generated inactive ADGRE1 (light gray) in side view **(A)** and extracellular view **(B)** reveals that β-arrestin 1-coupled ADGRE1 adopts an active conformation, characterized by an outward movement of the intracellular end of the TM6 helix. **(C-D)** Structural comparison of the β-arrestin 1 in the ADGRE1-bound state with the inactive state (C, PDB: 1G4M) and active state (D, PDB: 4JQI) shows that β-arrestin 1 in ADGRE1-coupled state adopts an active conformation, characterized by an interdomain twist of approximately 15 degrees. **(E)** Superposition of ADGRE1-βarr1 complex with previously solved receptor-arrestin complex structures, including NTSR1-βarr1 (PDB:6UP7), M2R-βarr1 (PDB: 6U1N), β1AR-βarr1 (PDB:6TKO), CB1-βarr1 (PDB:8WU1), V2R-βarr1 (PDB: 7R0J), and 5-HT2b-βarr1 (PDB: 7SRS). **(F)** Illustration of the angular relationships between receptors and β-arrestin1 cross different complexes, including ADGRE1-βarr1, β1AR-βarr1 (PDB:6TKO), CB1-βarr1 (PDB:8WU1), V2R-βarr1 (PDB: 7R0J), NTSR1-βarr1 (PDB:6UP7), 5-HT2b-βarr1 (PDB: 7SRS), and M2R-βarr1 (PDB: 6U1N).

**Supplemental Figure 9: Metadynamics simulations of ADGRE1-arrestin1 complex. (A)**

Free energy surface (FES) obtained from metadynamics, llustrating the energetic landscape of ADGRE1-arrestin dimerization. **(B)**Two-dimensional energy profile depicting the energy changes along key reaction coordinates. **(C)** Residue contact frequency analysis between the two β-arrestin1 protomers at the dimerization interface in the ADGRE1-arrestin1 dimerization interface.

### RESOURCE AVAILABILITY

#### Lead Contact

Further information and requests for resources and reagents should be directed to and will be fulfilled by the corresponding author, Sudarshan Rajagopal.

### Materials Availability

All plasmids generated in this study will be distributed upon request

### EXPERIMENTAL MODEL AND SUBJECT DETAILS

HEK 293T cells, including a β-Arrestin 1/2 knockout (KO) variant, were cultured in Dulbecco’s Modified Eagle Medium (DMEM) with 10% fetal bovine serum (FBS) and 1% antibiotic-antimycotic solution from Gibco, under conditions of 37°C and 5% CO2 humidity. The β-Arrestin 1/2 KO cells, created through CRISPR/Cas9 genome editing, were authenticated by immunoblot analysis and obtained from Dr. Asuka Inoue.

For experiments requiring confocal microscopy, both HEK293T and β-Arrestin 1/2 KO cells were seeded on 35-mm glass-bottom dishes pre-coated with either poly-D-lysine or rat tail collagen, aiming for a confluence between 40-70%. In the case of 96-well plate assays, cell density was adjusted to 70,000-100,000 HEK293T cells per well.

Transient transfections were performed using OPTI-MEM and Polyethyleneimine (PEI) at a PEI to DNA mass ratio of 3:1. Cells designated for confocal microscopy analysis were prepared and imaged after 16-24 hours post-transfection, adhering to the same timeline for BRET and split nanoluciferase assays.

## METHOD DETAILS

### Generation of Constructs

Constructs were developed using a modified overlap cloning technique^90^. Flexible linkers, consisting of glycine-serine repeats (GGGGS), with lengths ranging from 18 to 33 amino acids, were inserted between the coding sequences for fluorescent proteins or luciferases and those of receptors, transducers, biosensors, or adaptor proteins. The creation of SmBiT-β-Arrestin IP6, AP2 and Clathrin mutants utilized a previously published overlap cloning strategy with smBiT. The 2xFYVE-mKO construct was created by integrating Cyto-mKO into the 2xFYVE-LgBiT vector via overlap cloning.

Primers for overlap cloning can be found in **Table S2.**

### Generation of **β**-Arrestin Split-GFP

β-Arrestin Split GFP constructs were derived from pCDNA3.1-GFP1-10 (Addgene #70219) and peGFP-GFP11-Actin (Addgene #181966) and cloned into either SmBiT-β-Arrestin or β-Arrestin-SmBiT.

### Mutagenesis of **β**-Arrestin IP_6_ and IDR mutants

Mutation generation was carried out using the QuikChange site-directed mutagenesis kit (Agilent). The mutations produced included ΔIP_6_-N, ΔIP_6_-C, ΔIP_6_-N/C, ΔIP_6_-NT, ΔIP_6_-NT/C, ΔIDR, and ΔCT.

### Opto **β**-arrestin Constructs

Opto β-arrestin constructs were designed from previously published work.^47^ pHR-mCh-Cry2WT (Addgene 101221) and pHR-FUSN-mch-Cry2WT (Addgene #101223) were cloned into the pcDNA backbone. Overlap cloning was then used to generate Opto-β-Arrestin 1, Opto-β-Arrestin 2, Opto-β-Arrestin 1 IDR, Opto-β-Arrestin 2 IDR

### Generation of Constructs for ADGRE1-**β**-Arrestin1-scFv30 Complex

For structure determination of the ADGRE1-β-Arrestin1-scFv30 complex, the full-length mouse ADGRE1 gene (UniProt ID: Q14246) was cloned into the pFastBac1 vector. The endogenous signal peptide of ADGRE1 was replaced with an influenza hemagglutinin (HA) signal peptide, and an N-terminal FLAG tag (DYKDDDDK) followed by a thermostabilized cytochrome b562RIL (BRIL) fusion were introduced to enhance expression, stability, and facilitate purification. To stabilize the β-arrestin 1 complex, we used a truncated version of Bos taurus β-arrestin 1 (residues 1-376) with seven cysteine mutations (C59A, C125S, C140I, C150V, C242V, C251V, and C269S) to minimize surface-exposed cysteine residues, as previously reported to improve protein yield, stability and reduce aggregation. An additional R169E mutation was introduced to maintain β-arrestin 1 in its active state. A single-chain fragment variable 30 (scFv30) was fused to the C-terminus of β-arrestin 1 using a linker (KKNIAFLLASMFVFSIATNAYA) to stabilize of the complex, and the construct was cloned into the pFastBac1 vector with a C-terminal hexa-histidine tag. Full-length wild-type human GRK2 was also cloned into the pFastBac1 vector with a C-terminal hexa-histidine tag. All mutations were introduced using the QuikChange Site-Directed Mutagenesis kit (Thermo Fisher Scientifc) following the manufacture’s instructions. All constructs and mutations were verified by DNA sequencing.

### Protein expression and purification of ADGRE1-β-arrestin1-scFv30 complex

Following successful construct generation, recombinant baculoviruses for each component were generated using the Bac-to-Bac system (Invitrogen) in *Spodoptera frugiperda* (Sf9) cells, which were routinely tested and confirmed to be free from mycoplasma contamination. Baculovirus stock (P1) was harvested and subsequently used to infect Sf9 cells. For protein expression, Sf9 cells were cultured in ESF921 serum-free medium to a density of 2.5 × 10 cells per mL, followed by co-infection with recombinant baculoviruses encoding ADGRE1, β-arrestin1, and GRK2 at a multiplicity of infection (MOI) ratio of 1:1:1. Infected cells were incubated at 27°C with shaking at 110 rpm for 48 hours before being harvested by centrifugation, flash-frozen, and stored at-80°C until use.

For complex formation and purification, frozen cell pellets were thawed and resuspended in lysis buffer containing 20 mM HEPES (pH 7.5), 100 mM NaCl, and 3 mM MgCl₂, supplemented with cOmplete Protease Inhibitor Cocktail (Roche). To stabilize the complex, 10 mM iodoacetic acid (pH 8.0), 1 mM sodium orthovanadate, 1 mM sodium pyrophosphate, and 10 mM NaF were added, and the suspension was incubated at room temperature for 30 minutes. Membrane solubilization was carried out by adding 0.5% (w/v) lauryl maltose neopentyl glycol (LMNG; Anatrace) and 0.05% (w/v) cholesteryl hemisuccinate (CHS) for 2 hours at 4°C with gentle agitation. Insoluble debris was removed by ultracentrifugation at 65,000 × g for 30 minutes, and the solubilized receptor-arrestin complex was captured by batch binding to M1 anti-FLAG affinity resin in the presence of 5 mM CaCl₂. The resin was packed into a gravity-flow column and washed with 30 column volumes of washing buffer containing 20 mM HEPES (pH 7.5), 100 mM NaCl, 3 mM MgCl₂, 5 mM CaCl₂, 0.01% (w/v) LMNG, and 0.001% (w/v) CHS. The M1-resion bound proteins were further eluted with washing buffer supplemented with 10 mM EGTA and 0.2 mg/mL FLAG peptide. The eluted ADGRE1-β-arrestin1-scFv30 complex was concentrated using a 100-kDa molecular weight cutoff (MWCO) Amicon Ultra centrifugal filter (Millipore) and subjected to size-exclusion chromatography (SEC) on a Superose 6 Increase 10/300 column (Cytiva) pre-equilibrated in SEC buffer containing 20 mM HEPES (pH 7.5), 100 mM NaCl, 0.00075% (w/v) LMNG, 0.0001% (w/v) CHS, and 0.00025% (w/v) glyco-diosgenin (GDN; Anatrace). The fractions corresponding to the ADGRE1-β-arrestin1-scFv30 complex were pooled, and the purity, as well as the stoichiometry of complex components, was assessed by SDS-PAGE. The final purified complex was concentrated to approximately 1.2 mg/mL for cryo-EM sample preparation.

### Cryo-grid preparation and data acquisition

For cryo-EM grid preparation, 3.0 μl of the purified ADGRE1-FL-β-arrestin1-scFv30 complex (1.2 mg/ml) was applied onto a glow-discharged holey carbon grid (Quantifoil R1.2/1.3, 300 mesh) and vitrified in liquid ethane using a Vitrobot Mark IV (Thermo Fisher Scientific) under 4°C and 100% humidity, with parameters set to 3 s blot time, blot force of 1, and 10 s wait time. Verified grids were stored in liquid nitrogen until use. Cryo-EM data acquisition was performed on a Cs-corrected Titan Krios G4 microscope (300 kV) equipped with a Selectris X imaging filter and Falcon 4 camera. Images were collected at 81,000× magnification (0.92 Å/pixel) using SerialEM with automated low-dose acquisition. Movie stacks were recorded at a defocus range of-1.0 to-2.0 μm, with a total accumulated dose of 60 e⁻/Å² distributed over 32 frames per micrograph.

### BRET and split Nano Luciferase assays

For comprehensive BRET and split luciferase assays, HEK293 cells were cultured in 6-well plates and transiently transfected utilizing Polyethylenimine (PEI) as outlined previously. The β-Arrestin association was analyzed using split NanoLuc components (i.e., SmBiT and LgBiT) to explore various protein-protein interactions, including amino-amino (N-N), amino-carboxyl (N-C), and carboxyl-carboxyl (C-C) interfaces. Specifically, 500 ng of the receptor and 100 ng of each β-Arrestin construct were introduced into the cells. In the NanoBiT-BRET assay setup, 500 ng of the receptor and 1 μg of location-specific tagged-mKO (including Cyto-mKO, CAAX-mKO, AP2-mKO, 2xFYVE-mKO) were co-transfected with 200 ng of each NanoBiT β-Arrestin variant. Cyto-mKO and CAAX-mKO were used for the normalization of β-Arrestin association studies (e.g., N-C interactions). For β-Arrestin recruitment assays, cells received 500 ng of receptor-LgBiT and 200 ng of N-terminally tagged SmBiT-β-Arrestin. To ascertain β-Arrestin’s recruitment to specific cellular locales, 500 ng of the receptor and 1 μg of either CAAX-LgBiT or 2xFYVE-LgBiT were used. Endocytosis assays were performed by co-transfecting 200-500 ng of receptor-Rluc with 1-1.5 μg of 2xFYVE-Venus. For EKAR assays, 25 ng of the biosensor was paired with 1 μg of the receptor and 200 ng of β-Arrestin. For TRUPATH assay, 1:1:1:1 μg of receptor: Glll_q_: Gβ: G_Y_ were used^79,80^. On the following day (Day 2 post-transfection), cells underwent washing with phosphate-buffered saline (PBS), detachment via trypsinization, and seeding onto a Corning Costar 96-well clear bottom, white-walled plate at a density of 70-100,000 cells per well. The culture medium was replaced with clear minimal essential medium, enhanced with 2% FBS, 1% P/S, 10 mM HEPES, 1x GlutaMAX, and 1x Antibiotic-Antimycotic (Gibco). On Day 3, the culture medium was removed, and cells were incubated with 80 μL of 3 μM coelenterazine h in HBSS, further supplemented with 20 mM HEPES for 5 minutes. Prior to ligand addition in split luciferase assays, three baseline reads were recorded to assess basal luminescence, which was then normalized to vehicle control conditions and shown as % change in luminescence. Luminescence and BRET ratios were quantified using a BioTek Synergy Neo2 plate reader at 37°C. For BRET measurements, a 480 nm wavelength filter for the donor and a 530 nm or custom mKO 542 nm long-pass emission filter for the acceptor were employed. Net BRET was determined by subtracting the vehicle BRET ratio from the ligand-induced BRET ratio.

### Confocal Microscopy

HEK293 cells were seeded onto 35-mm dishes coated with poly-D-lysine and cultured until they reached 50-70% confluence. Following transfection using Polyethylenimine (PEI), cells were incubated for an additional 16-24 hours to ensure adequate expression of the transfected constructs. Cells were then washed with PBS and serum-starved for 1 hour to synchronize cellular responses. Prior to imaging, cells were treated for 5 minutes at 37°C with either 16HD or a control serum-free medium. Subsequent to this pretreatment, cells were stimulated with ligands: 10 μM Isoproterenol, 1 μM Angiotensin II (AngII), or 1 μM Vasopressin (AVP). After stimulation, cells were fixed using 4% paraformaldehyde supplemented with Hoechst 33342 (1:1000, Thermo Fisher Scientific, #62249) for nuclear staining. Imaging was performed on a Zeiss 880 or 980 confocal microscope, utilizing appropriate laser lines for Hoechst 33342 (400 nm), GFP (480 nm), and mKO (548 nm).

### Live Cell Microscopy

For live-cell imaging, HEK293 cells were similarly prepared and transfected as described for confocal microscopy. After the serum starvation period, cells were placed in a live-cell chamber system equipped with a temperature stage at 37°C. All live cell imaging was performed using 63X objective. For optogenetic β-arrestin experiments, two laser wavelengths were used (488nm for Cry2 activation and 560nm for mCherry).

### Fluorescence recovery after photobleaching (FRAP)

FRAP was conducted using a Zeiss 980 confocal microscope with a 63X objective, leveraging a 488 nm laser for targeted bleaching of regions of interest (ROIs). The procedure aimed to observe fluorescence recovery within these ROIs over 3 minutes at specified intervals. Small circular ROIs were designated on either punctate structures or the diffuse cytosol, and bleaching was performed with the laser at 100% power to diminish fluorescence selectively. Following bleaching, fluorescence recovery was captured, allowing for the analysis of protein dynamics. Recovery data was processed using ImageJ for initial quantification. Subsequently, Microsoft Excel was utilized to normalize the fluorescence intensity data, setting the 5 pre-bleach values to 1 for a standardized baseline and the immediate post-bleach intensity to 0.

### Immunoblotting

Immunoblotting procedures were performed in accordance with previously established protocols. HEK293 cells were cultured in 6-well plates and transiently transfected with β-arrestin pcDNA constructs using Polyethyleneimine (PEI). Following a 24-hour post-transfection period, cells underwent serum starvation using Minimum Essential Medium (MEM). Subsequently, cells were cooled on ice, rinsed with ice-cold PBS, and lysed using a buffer containing protease inhibitors Phos-STOP (Roche) and cOmplete EDTA-free (Sigma). Lysates were agitated at 4°C for 45 minutes and then centrifuged at over 12,000g for 15 minutes at 4°C to remove insoluble debris. The resulting supernatant was processed further. Protein samples were separated on SDS-10% polyacrylamide gels and transferred onto nitrocellulose membranes for blotting. Primary antibodies targeting phospho-ERK (1:1000 dilution, Cell Signaling Technology, #9106) and total ERK (1:1000 dilution, Millipore Sigma, #06-182) were applied overnight to evaluate ERK activation. The A1-CT antibody, specific for β-arrestin isoforms, and alpha-tubulin (Sigma-Aldrich, #T6074) as a loading control were also used. Detection was facilitated by horseradish peroxidase-conjugated secondary antibodies (mouse anti-rabbit IgG or anti-mouse IgG) at a 1:3000 dilution. The detection of immune complexes on the membranes was achieved using SuperSignal enhanced chemiluminescent substrate (Thermo Fisher) and documented with imaging equipment.

### Acquisition of β-arrestin puncta

On day one, HEK293 cells were transfected in 6-well dishes and incubated for 24 hours. On day two 70,-100,000 cells were plated on poly-D-lysine coated plates (Thermo Scientific, 152037). On day three, cells were fixed with 4% PFA. After 30 minutes of fixation, cells were washed 1xPBS for 10 minutes x 3.

### Image processing and 3D reconstructions

The overall cryo-EM data processing workflow for the ADGRE1-β-arrestin1-scFv30 complex is summarized in Supplemental Figure 6. Raw cryo-EM movie stacks were first corrected for dose-weighted beam-induced motion using UCSF MotionCor2^91^, and the contrast transfer function (CTF) parameters were estimated using patch-based CTF estimation in CryoSPARC v4.4^92^. A total of 23, 906 movie stacks were imported into CryoSPARC, with each micrograph individually inspected to exclude images exhibiting crystalline ice contamination or other artifacts. For optimal particle picking, the Topaz deep-learning neural networks algorithm within CryoSPARC was employed. A total of 2,549,811 particles were extracted at an initial box size of 320 pixels, binned to 160 pixels, and subjected to 2D classification. From this dataset, 249,976 high-quality particle projections were selected and re-extracted at the full 320-pixel box size. These particles were then subjected to ab initio reconstruction and heterogeneous refinement with enforced C₂ symmetry for 3D classification. The best-resolved class, containing 140,588 particle projections, was further refined using homogeneous refinement followed by non-uniform refinement with C₂ symmetry, yielding the final density map at 3.58 Å global resolution.

### Model building and refinement

The density map for ADGRE1-β-arrestin1-scFv30 complex were sharpen with Deep EMhancer prior to model building. The initial ADGRE1 (GPR133) model was generated by AlphaFold2, while β-arrestin1 and scFv30 were derived from the [xxxx, PDB] structure. The initial model of the ADGRE1-β-arrestin1-scFv30 complex was built by docking these components into the maps using Chimera, followed by iteratively adjusted via manual rebuilding in Coot and real-space refinement in Phenix. Final refinement statistics are listed in Table S1. Structural figures were prepared using ChimeraX^93^ or Pymol.

### Metadynamics of ADGRE1-**β**-arrestin1 complex

To determine whether the N-N orientation presented in ADGRE1-β-arrestin1 complex facilitates the direct interactions β-arrestins, we performed metadynamics (MetaD) simulations^94^ on ADGRE1-β-arrestin 1 complex. First, missing residues and side chains in the cryo-EM structure were completed using Maestro. The simulation parameters for the complex system were then generated using CHARMM-GUI^95^, and the CHARMM36m force field^96^ was applied. The complex was embedded into a pre-equilibrated 1-palmitoyl-2-oleoyl-sn-glycero-3-phosphocholine (POPC) lipid bilayer, solvated using the TIP3P water model, and neutralized with 0.15 M NaCl. The orientation of the protein within the membrane was optimized based on the Orientations of Proteins in Membranes (OPM) database^97^. Simulations were performed using GROMACS 2019.6^98^, starting with energy minimization (10,000 steps), combining 5,000 steps of the steepest descent method and 5,000 steps of the conjugate gradient method. This was followed by an NVT ensemble phase, where the system was heated from 0 K to 310 K over 10,000 ps. Subsequently, an NPT ensemble simulation was conducted at 1 atm for 10,000 ps, with a harmonic restraint of 10.0 kcal·mol⁻¹·Å⁻² applied to the system.Two collective variables (CV1 and CV2) were defined to describe the RMSD changes between interacting arrestin monomers, capturing the conformational evolution. Enhanced sampling was carried out using GROMACS 2019.6 and PLUMED 2.7.1^99^, with HILLS added every 500 steps, an initial Gaussian height of 10 kJ·mol⁻¹, a Gaussian width of 1,000 kJ·mol⁻¹, and a bias factor of 15.

## QUANTIFICATION AND STATISTICAL ANALYSIS

### Quantification of cellular puncta

For analysis, Image Xpress Pico Automated Cell Imagining System (Molecular Device) was used. Thresholds or size and image intensity were made to negative controls. Images were captured at 20x and ∼5000 cells were analyzed/per well. Puncta was normalized to each cell with the nuclear marker.

### Image Analysis

Confocal Images were visualized using ImageJ. All Image adjustments are identical and consistent.

#### Statistics and reproducibility

Data was analyzed in Excel and graphed in Prism 10 (GraphPad, San Diego, CA). Dose-response curves were fitted to log agonist versus stimulus with three parameters with the minimum baseline corrected to zero. Statistical tests were performed using a two-way ANOVA when comparing different β-arrestin mutants in time response assays AUC. Further details of statistical analysis and replicates can be found in the figure legend. Crucial plate-based experiments were independently replicated by at least two different investigators whenever feasible.

## Notes

### Competing Interest Statement

The authors have declared no competing interest.

### Summary of Updates

The original submission had blurry figures due to a PDF conversion issue. All figures have been updated

