## Supplementary figures and images for "β-Arrestin Condensates Regulate G Protein-Coupled Receptor Function"

### Supplementary Materials

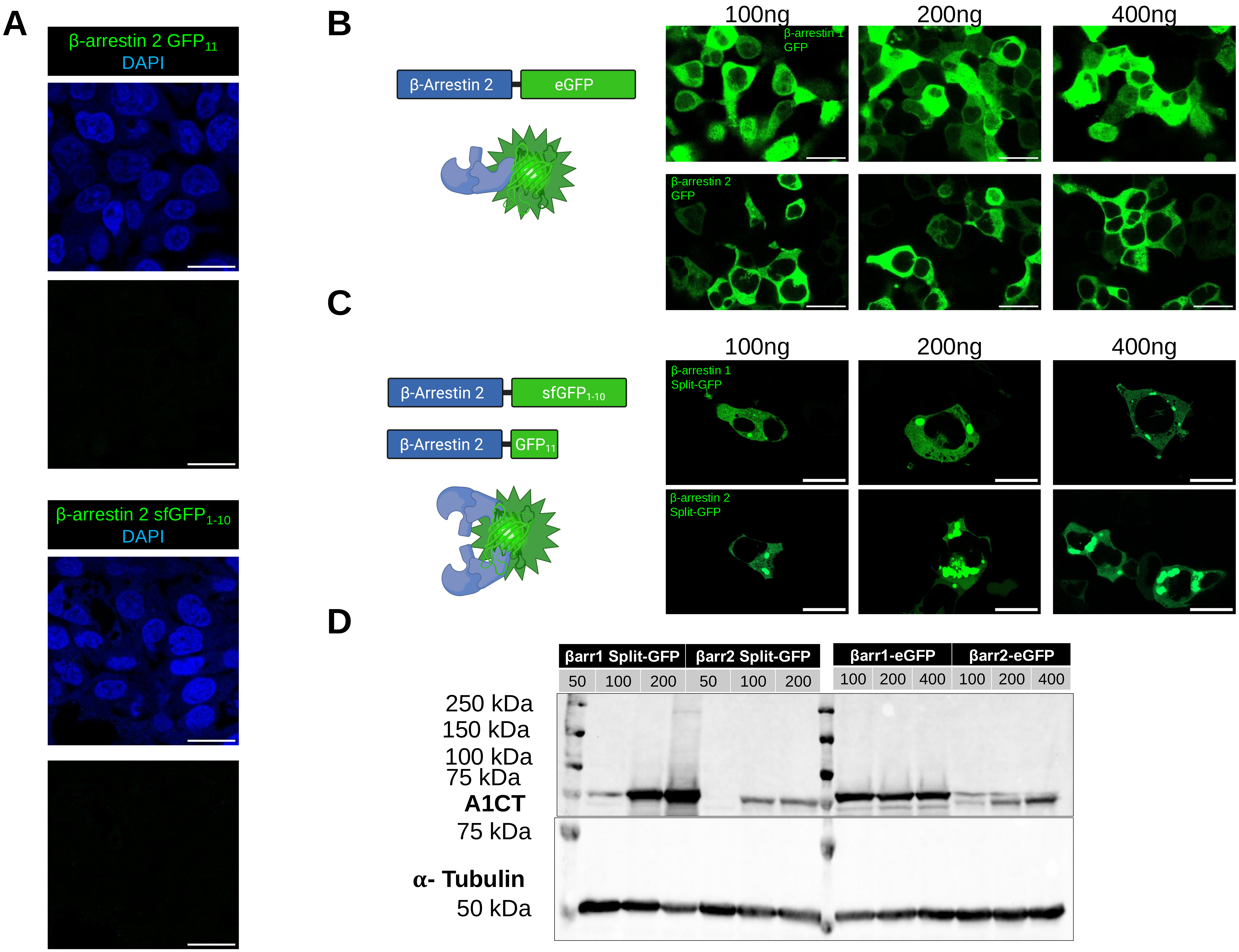

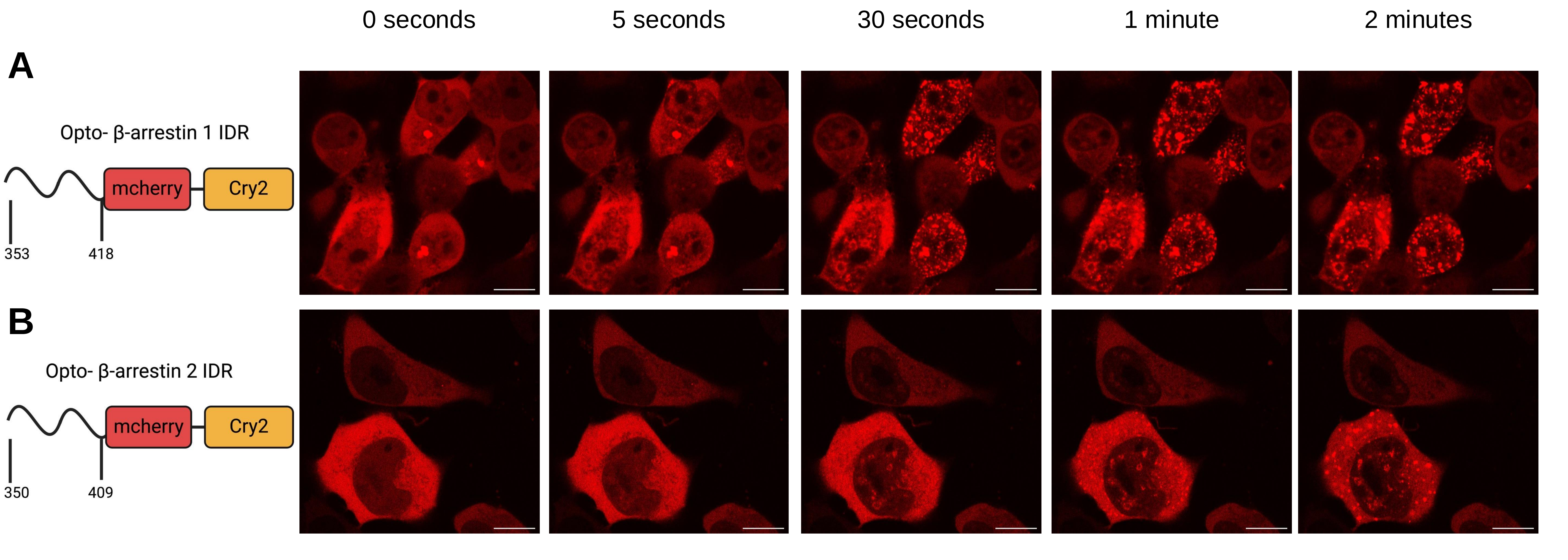

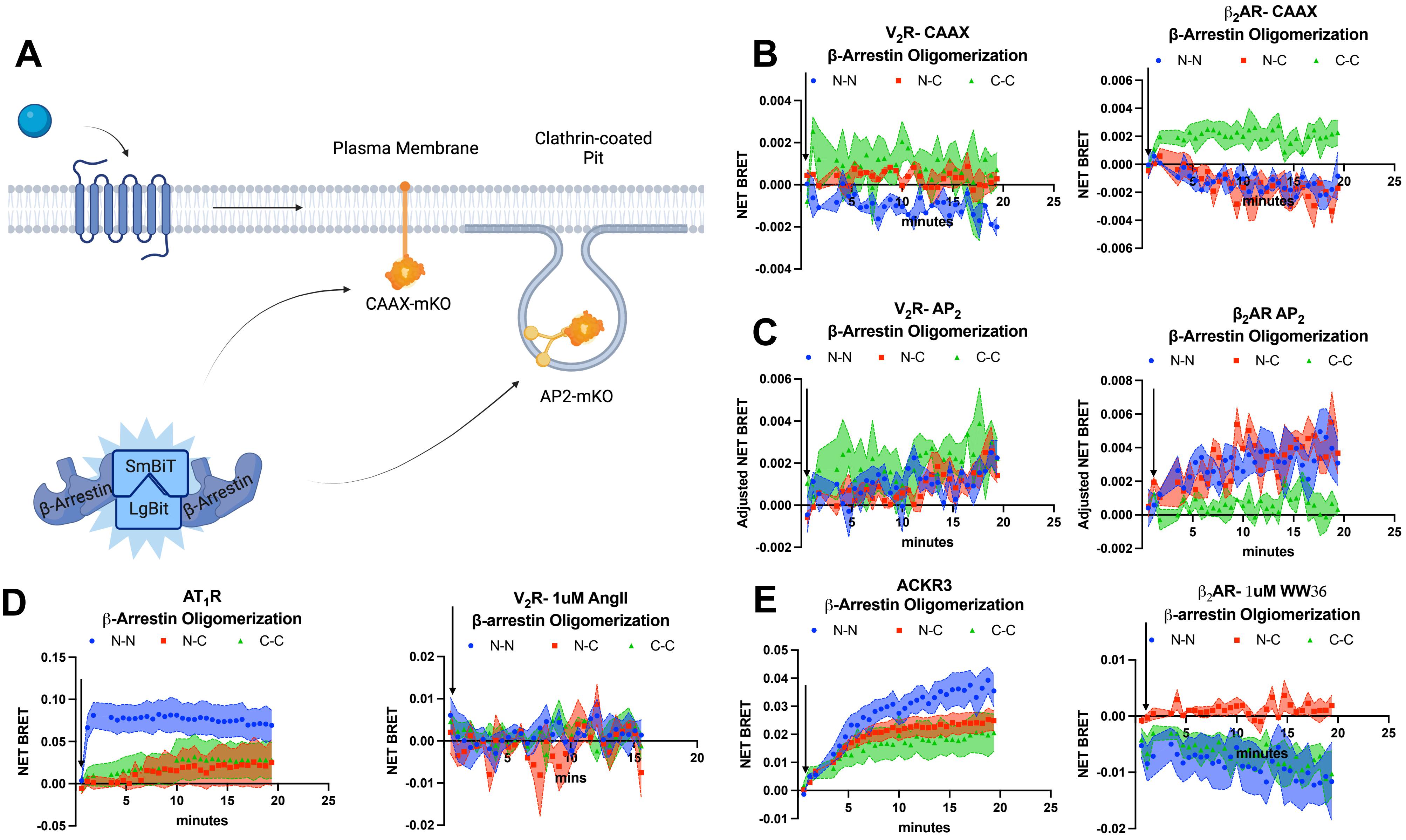

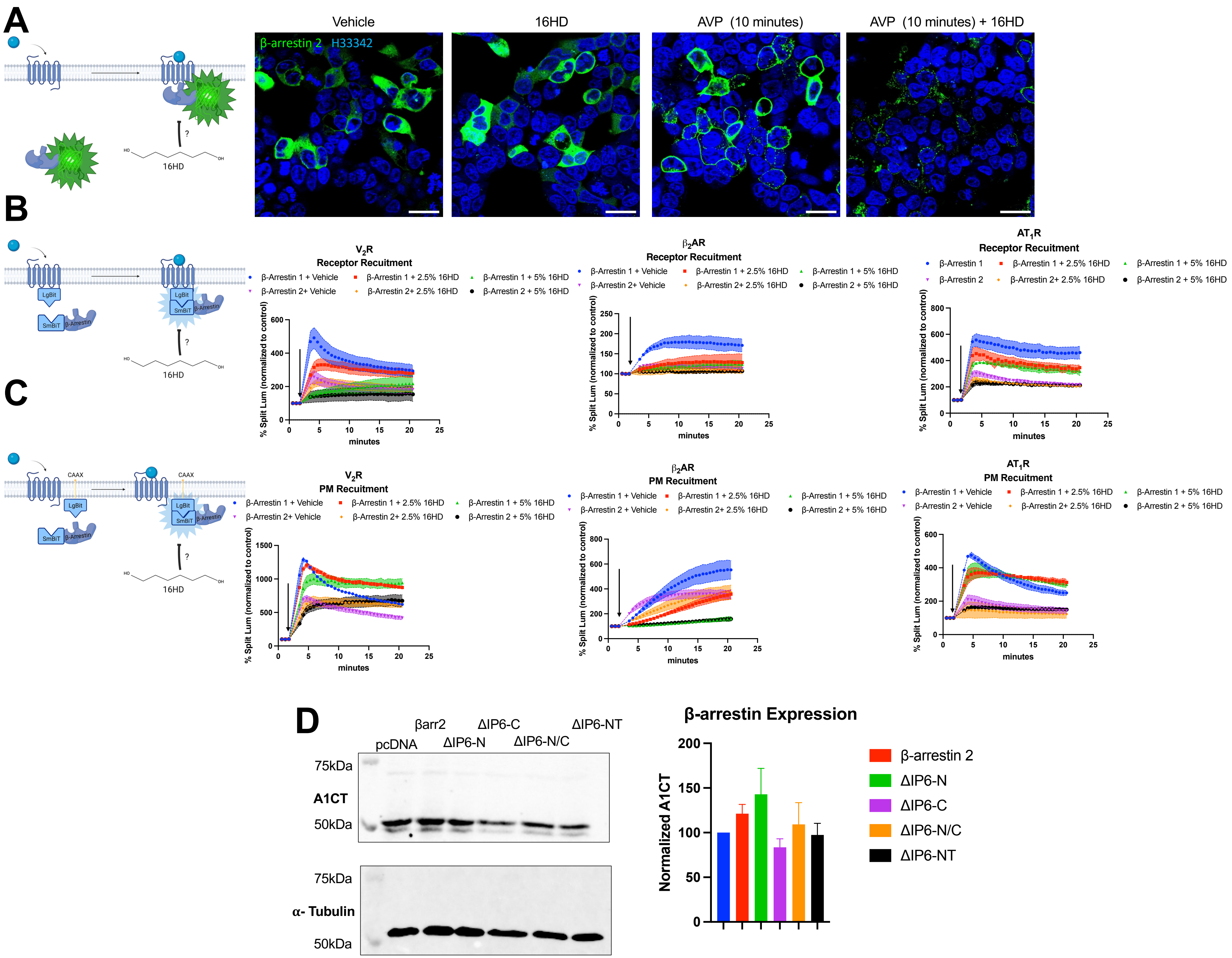

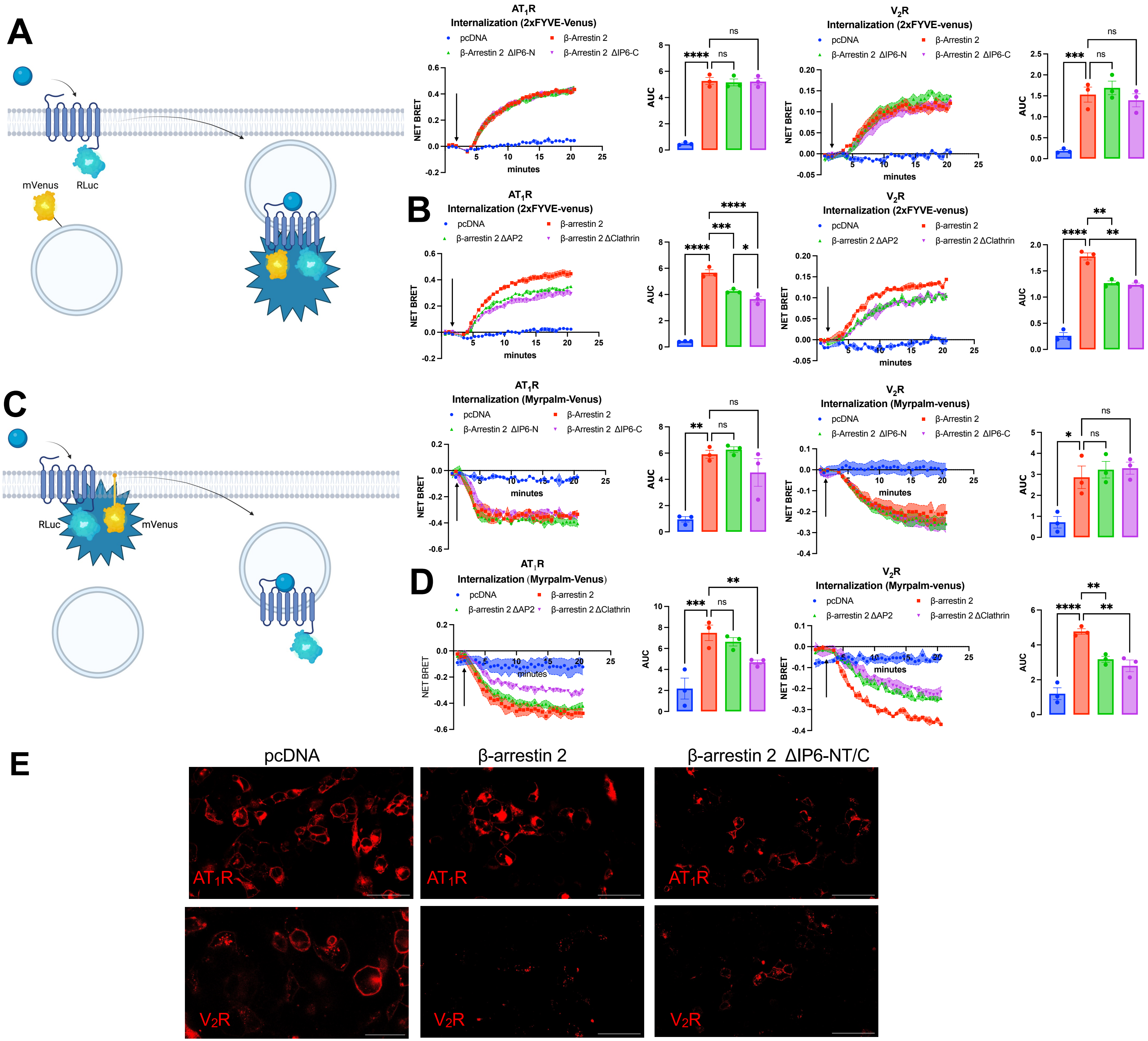

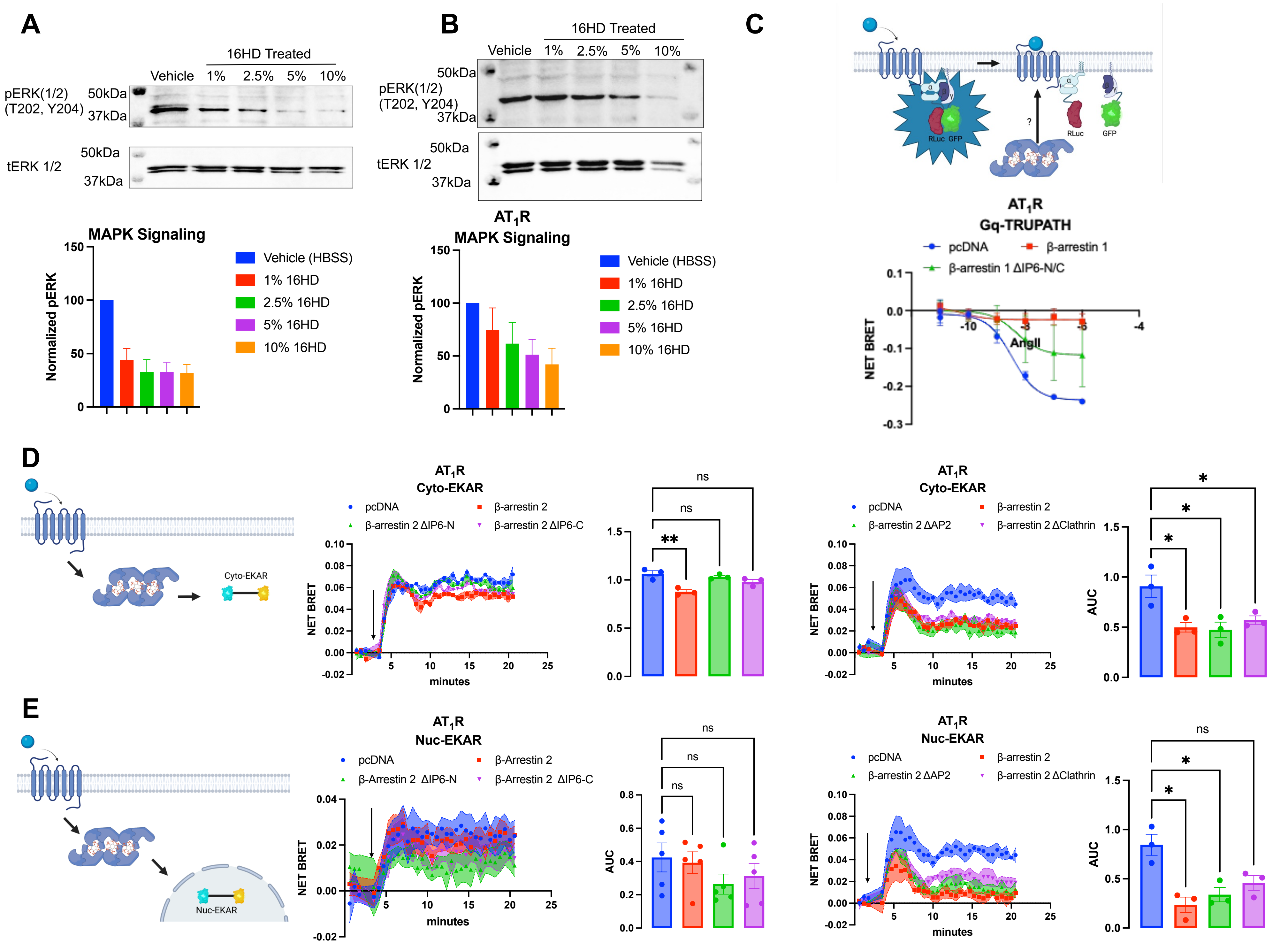

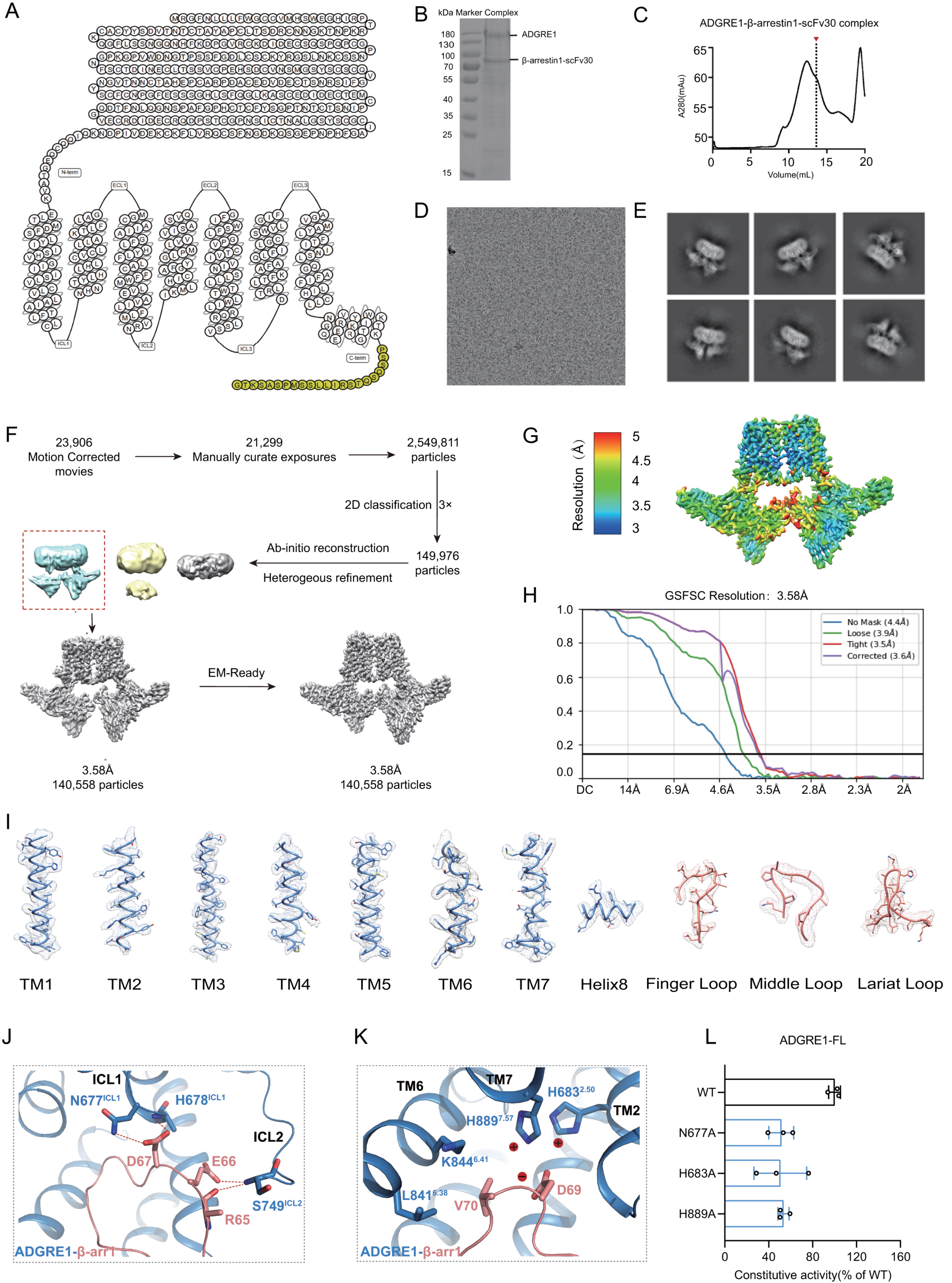

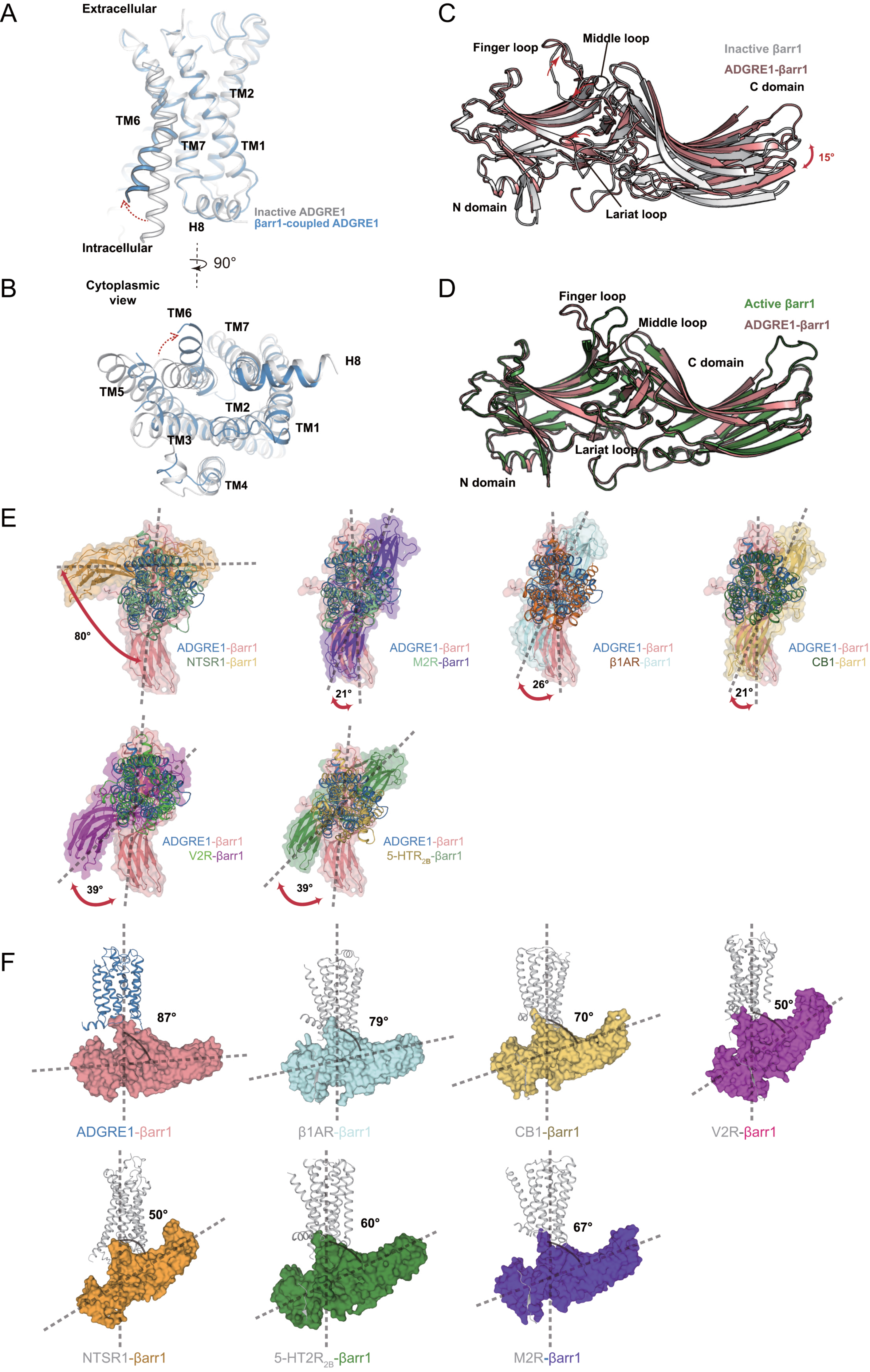

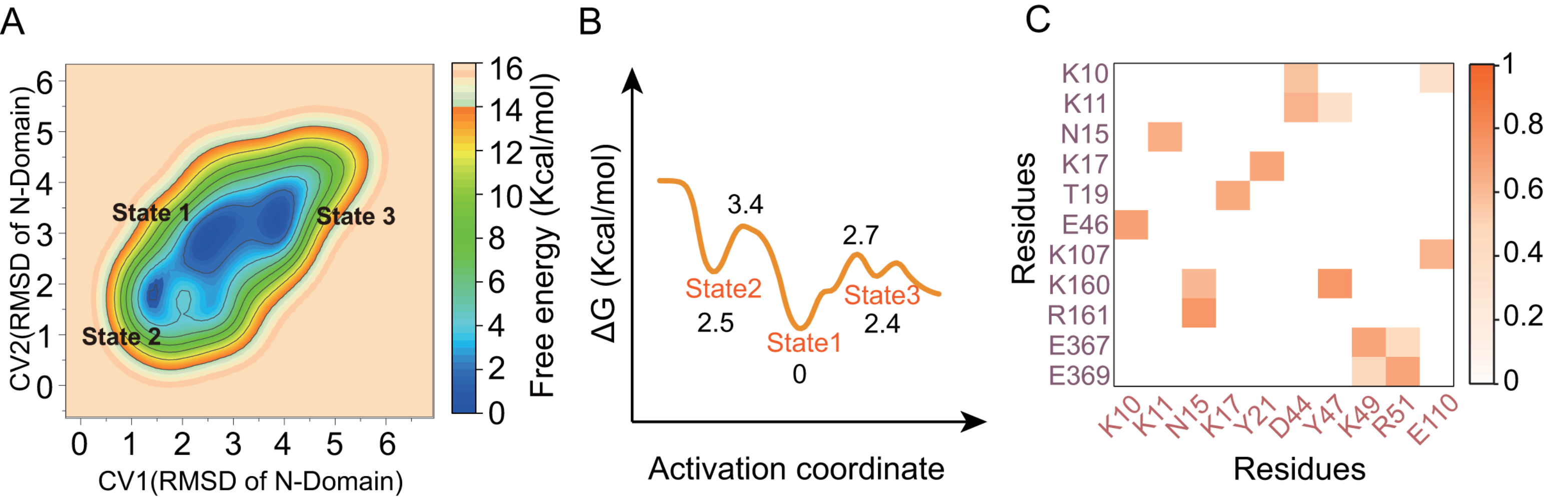
